# Different inhibition of Nrf2 by two Keap1 isoforms α and β to shape malignant behaviour of human hepatocellular carcinoma

**DOI:** 10.1101/2022.07.15.500244

**Authors:** Feilong Chen, Mei Xiao, Jing Feng, Reziyamu Wufur, Keli Liu, Shaofan Hu, Yiguo Zhang

**Author notes:** Correspondence: to YZ.

## Abstract

Nrf2 (nuclear factor E2-related factor 2, encoded by *Nfe2l2*) acts as a master transcriptional regulator in mediating antioxidant, detoxification and cytoprotective responses against oxidative, electrophilic and metabolic stress, but also plays a crucial role in cancer metabolism and multiple oncogenic pathways, whereas the redox sensor Keap1 functions as a predominant inhibitor of Nrf2 and hence changes in its expression abundance directly affect the Nrf2 stability and transcriptional activity. However, nuanced functional isoforms of Keap1α and β have rarely been identified to date. Herein, we have established four distinct cell models stably expressing *Keap1^-/-^, Keap1β* (*Keap1^Δ1-31^*), *Keap1-Restored* and *Keap1α-Restored*, aiming to gain a better understanding of similarities and differences of two Keap1 isoforms between their distinct regulatory profiles. Our experimental evidence revealed that although Keap1 and its isoforms are still localized in the cytoplasmic compartments, they elicited differential inhibitory effects on Nrf2 and its target HO-1. Furtherly, transcriptome sequencing unraveled that they possess similar but different functions. Such functions were further determined by multiple experiments *in vivo* (i.e. subcutaneous tumour formation in nude mice) and *in vitro* (e.g., cell cloning, infection, migration, wound healing, cell cycle, apoptosis, CAT enzymatic activity and intracellular GSH levels). Of note, the results obtained from tumorigenesis experiments in xenograft model mice were verified based on the prominent changes in the PTEN signaling to the PI3K-AKT-mTOR pathways, in addition to substantially aberrant expression patterns of those typical genes involved in the EMT (epithelial-mesenchymal transition), cell cycle and apoptosis.

## 1. Introduction

Keap1 (Kelch-like ECH-associated protein 1) was originally isolated by a yeast two-hybrid system[1]. The full-length Keap1 protein is composed of 624 amino acids (aa) with a molecular weight of approximately 69 kDa [2] and its gene contains six exons and five introns located on human chromosome 19p13.2 [3]. Nevertheless, there have been few reports on this protein isoforms. The functions of Keap1 protein isoforms (Keap1a and Keap1b) have been reported in zebrafish[4], and a novel natural mutant of Keap1 (Keap1^ΔC^) was reported [3]. This mutant Keap1^ΔC^ lacks most of the C-terminal Kelch/DGR domain of Keap1 that enables physical interaction with the Neh2 domain of Nrf2 (nuclear factor-erythroid 2-related factor 2, encoded by *Nfe2l2*), which is a key transcription factor of the cap’n’collar (CNC) basic-region leucine zipper (bZIP) family. Just due to this direct interaction, Keap1 has been identified as a master inhibitor of Nrf2, and its high or low expression levels in human lung, breast, liver and other somatic cancers hence causes changes in endogenous Nrf2 abundances [5,6]. The Nrf2-interacting Kelch/DGR domain of Keap1 is composed of six repeating Kelch sequences [7], whilst the Keap1-binding Neh2 domain is located at the N-terminus of Nrf2 and contains two highly conserved motifs comprising the ^29^DLG^31^ and ^79^ETGE^82^ sequences, respectively. Keap1 functions as a master inhibitory protein of Nrf2 by segregating the CNC-bZIP factor in the cytoplasm and then targeting this protein to the ubiquitin-mediated proteasomal degradation system [1,8,9].

As one of the most important members of the CNC-bZIP family[10,11], Nrf2 plays a crucial role in the management of cellular response to oxidative and electrophilic stress. Although Nrf2 originated in foreign organisms as a master regulator in response to oxidative or metabolic stress, it has been implicated in metabolism, as well as multiple oncogenic pathways in cancer cells[12]. Further studies have demonstrated the physiological importance of such Nrf2-dependent cytoprotective responses in *Nrf2*-deficient mice[13], rats [14] and zebrafish[15]. The unstimulated Nrf2 protein remained predominantly in the cytoplasm and directly bound to Keap1, allowing it to be targeted to the ubiquitin-mediated degradation pathway [16,17]. The protein degradation of Nrf2 is promoted by Keap1 through the ubiquitin ligase Cullin3 to the proteasome-mediated proteolysis [18,19]. Once Nrf2 is activated, it is separated from Keap1[20]. The activated Nrf2 is translocated into the nucleus, and bound to antioxidant response elements (AREs) in its cognate target gene promoter regions by its functional heterodimer with small MAF and other bZIP transcription factors [18]. Consequently, a large number of Nrf2-dependent antioxidant genes and detoxification enzymes are transcriptionally activated and expressed in order to meet the needs for physiological or biological functions. Nrf2-targets include HO-1 (heme oxygenase-1), GSR (glutathione reductase), NQO1 (NAD(P)H quinone oxidoreductase-1), GCLC (γ-glutamyl cysteine ligase catalytic subunit), and GCLM (γ-glutamate-cysteine ligase modifier subunit), amongst others [18,20].

Herein, we found that Keap1 has two main isoforms, which are yielded by alternative transcriptional translations and differentially expressed in different cells and tissues. For descriptive convenience, they were named Keap1α (larger molecular weight) and Keap1β (smaller molecular weight). To explore the functional differences between Keap1α and Keap1β, we constructed *Keap1^-/-^* and *Keap1β* (*Keap1^Δ1-31^*) cell lines, and the stable expression of Keap1 and Keap1α was restored by the lentiviral system into the *Keap1^-/-^* cell line. The results revealed that when Keap1 is knocked out, the cell proliferation and migration, and its tumour xenografts are reduced. Such negative effects were, however, reversed to varying degrees by restoring Keap1 and Keap1α in *Keap1^-/-^* cells. Then, a series of comparative experiments (subcutaneous tumour formation in nude mice, cell cloning, infection, migration, the CAT enzymatic activity, intracellular GSH levels, wound healing, EMT, cell cycle progression and apoptosis) were employed, so as to verify the similarities and differences in their regulatory functions of between Keap1α and Keap1β. Furthermore, transcriptome sequencing analysis unraveled that Keap1, Keap1α and Keap1β are involved in distinct signaling pathways.

## 2. Materials and methods

### 2.1. Cells, Cell Culture and Transfection

HepG2 and COS-1 cell lines used in this study were purchased from the Cell Bank of Type Culture Collection of Chinese Academy of Sciences (Shanghai, China). Both *Keap1^-/-^* and *Keap1β*(*Keap1^Δ1-31^*) cell lines were created by CRISPR/Cas9-guided gene editing technology. The *Keap1-Restored* and *Keap1α-Restored* cell lines were created by stably expressing the respective proteins using the lentivirus technology. These cell lines were treated according to the relevant experimental requirements. Transfection experiments was performed using the Lipofectamine 3000 Transfection Kit (Invitrogen), including one or more indicated plasmids being tranfected into cells, according to the manufacture’s instruction.

### 2.2. Chemicals and Antibodies

All chemicals were of the best quality commercially available. The protein primary antibody against Nrf1 was made in our own group [21]. Flag (DYKDDDDK) monoclonal antibody was purchased from Invitrogen. The secondary antibodies and β-actin were purchased from ZSGB-BIO (Beijing, China). All other main antibodies used in this study were listed in Table 1.

**Table 1.** Main antibody information in the text

| Antibodies | Identifier | Source |
| --- | --- | --- |
| Nrf1 | N/A | Zhang laboratory[21] |
| Nrf2 | ab62352 | Abcam |
| HO-1 | ab68477 | Abcam |
| NQO-1 | Ab80588 | Abcam |
| Flag | DYKDDDDK Tag Monoclonal Antibody, FG4R | Invitrogen |
| α-Tubulin | ab52866 | Abcam |
| Histone 3 | ab192985 | Abcam |
| PTEN | ab267787 | Abcam |
| PI3K | ab245781 | Abcam |
| AKT1 | ab32505 | Abcam |
| mTOR | ab134903 | Abcam |
| SNAI1 | A00716-2 | BioSynthesis |
| SNAI2 | PB9439 | BOSTER |
| CDH2 | ab76011 | Abcam |
| FN1 | BA1772 | BOSTER |
| MMP9 | BA0573 | BOSTER |
| Vimentin | Ab92547 | Abcam |
| Fluorescent secondary antibody(rabbit) | ZF-0511, Alexa Fluor 488 | ZSGB-BIO |
| Fluorescent secondary antibody(mouse) | ZF-0513, Alexa Fluor 594 | ZSGB-BIO |
| β-actin | TA-09 | ZSGB-BIO |

### 2.3. Expression Constructs

The human full-length Keap1 and Nrf2 cDNA sequences were subcloned into a pcDNA3 or 3XFlag vector, respectively. The Keap1α-expressing plasmid was obtained by mutating the N-terminal 2^nd^ translation starting codon ATG of the full-length Keap1 (i.e., located at the 32^nd^ residue position). The first N-terminal 31 amino acids (the first 93 bases) of Keap1 were truncated to yield the mutant Keap1β (i.e., *Keap1^Δ1-31^*).

### 2.4. Stable expression of Keap1^-/-^, Keepβ1(Keap1^Δ1-31^), Keap1-Restored and Keap1a-Restored

In HepG2 cells, CRISPR/Cas9-mediated genomic editing technology was used for knockout of Keap1 to yield two distinct cell lines *Keap1^-/-^* and *Keap1^-/-^ Keap1β^Δ1-31^*), which were constructed and identified by DNA sequencing (Figures 1A, 1B, and S1A-D). Both Keap1- and Keap1α-expressing plasmids were cloned into the pLVX-IRES-ZsGreen-puro vector for lentivirus packaging, according to previously-reported methods [22]. Then, *Keap1^-/-^* cells were infected with Keap1- or Keap1α-expressing lentiviral plasmids, and the stably expressing cell lines (i.e., *Keap1-Restored* and *Keap1α-Restored*) were constructed and identified by quantitative real-time PCR (q-PCR) and Western blotting (WB) experiments (Figure 1C). These experimental cell lines were positively selected by allowing them to be cultured in the puromycin-containing media.

### 2.5. RNA Isolation and Quantitative Real-Time PCR (q-PCR)

Total RNAs were prepared from cells using the RNA Simple Total RNA Kit (DP419, Tiangen Biotech Co., Ltd., Beijing, China), and reverse transcription was performed using the Revert Aid First Strand cDNA Synthesis Kit from Thermo. Then real-time quantitative PCR was carried out as previously described.

### 2.6. Analysis for the Nucleocytoplasmic Fractions

Experimental cells (4.0 × 10^5^) were grown on 6-cm cell culture plates for 48 h. The subcellular fractionation was conducted according to the instruction of nucleocytoplasmic separation kit ( Sigma, N-3408-200ML). The nucleocytoplasmic fractions were then determined by Western blotting with the relevant antibodies.

### 2.7. Cell Migration Analysis

Transwell assays were performed in modified Boyden chambers (8 μm, 24-well insert; Transwell; Corning Inc. Lowell, MA, USA). Briefly, after the growing cells reached 75%-85% confluency, they were starved in a serum-free medium for 12 h. Then a fresh medium containing foetal bovine serum was added into the lower chamber, whilst experimental cells (5 × 10^3^ suspended in 500 μL of a serum-free medium) were added into the upper chamber. The cells were cultured at 37 °C for additional 24 h, before they were fixed with a pre-cooled paraformaldehyde fixative (Serbicebio) for 30 min and stained with crystal violet (Beyotime) for 30 min at room temperature. All the above operations required for interval washing with phosphate-buffered saline (PBS) three times. The resultant cells were photographed and then counted as shown graphically.

### 2.8. Colony-formation Analysis

Equal numbers (1 ×10^3^) of experimental cells (*Keap1^+/+^, Keap1^-/-^, Keap1-Restored, Keap1α-Restored* and *Keap1β*(*Keap1^Δ1-31^*)) were allowed to grow for 10 days. The similar experiments were independently repeated three times. The resulting colonies were fixed with a precooled paraformaldehyde fixative (Serbicebio) for 30 min and stained with crystal violet (Beyotime) for 30 min at room temperature. All the above operations required for interval washing with PBS three times, and the cell clones were photographed and counted.

### 2.9. Wound-healing Analysis

After experimental cells (4 × 10^5^) were grown in 6-well plates to reach 80-85% confluency, they were starved in a serum-free medium for 12 hours. The cell monolayer was wounded with a 20-μL pipette tip and removed by washing with PBS. The migrating distance of the remaining cells was measured under microscope at indicated time points (0, 24, 48 and 72 h).

### 2.10. Assays for both CAT Activity and GSH Levels

Experimental cells (4 × 10^5^) were grown on 6-cm cell culture plates for 24 h. After the cells were digested, they were collected into 1.5 ml centrifuge tubes. The activity of catalase (CAT) and intracellular reduced glutathione (GSH) levels were assayed by using their detection kits, respectively (from Solarbio, BC0205 and Nanjing Jiancheng Bioengineering Institute, A006-2-1). The above experiments were independently repeated three times.

### 2.11. Cell Cycle and Apoptosis Detection by Flow Cytometry

Experimental cells (4 × 10^5^) were grown on 6-cm cell culture plates for 24 h. Afterwards, the cells were starved in a serum-free medium for 12 h before being treated with 10 μmol/L BrdU for additional 12 h. The cells were fixed for 15 min with 100 μl of BD Cytofix/Cytoperm buffer (containing a mixture of paraformaldehyde as a fixative and saponin as a detergent) at room temperature and then permeabilized for 10 min with 100 μl of BD Cytoperm permeabilization buffer plus (containing foetal bovine serum as a staining enhancer) on ice. Thereafter, the cells were refixed and treated with 100 ml DNase (at a dose of 300 mg/ml in DPBS) for 60 min at 37 °C to expose the incorporated BrdU, followed by staining with FITC-conjugated anti-BrdU antibody for 1 h at room temperature. Subsequently, the cells were suspended in 20 ml of 7-amino-actinomycin D solution for 20 min for DNA staining and resuspended in 0.5 ml of additional staining buffer (1× DPBS containing 0.09% sodium azide and 3% heat-inactivated FBS) prior to cell cycle analysis by flow cytometry. Furtherly, similar experimental cells (6 × 10^5^) were also allowed to grow for 48 h in 6-cm dishes before being harvested for apoptosis analysis. The cells were pelleted by centrifuging at 500× *g* for 5 min and washed with PBS three times before being incubated for 15 min with 5 ml of Annexin V-FITC and 10 ml of propidium iodide (PI) in 195 ml of the binding buffer, followed by flow cytometry analysis of cell apoptosis. These resulting data were obtained using a low flow rate of 400 events per second and further analyzed by FlowJo 7.6.1 software[23].

### 2.12. Western Blotting Analysis

After experimental cells were washed with PBS three times, total lysates were subjected to protein extraction. Total protein extracts were immediately denatured at 100 °C for 10 min. The eluted proteins were separated by SDS-PAGE containing 8%, 10 % and 15% polyacrylamide, and then visualized by immunoblotting with different primary antibodies against Keap1, Nrf2, HO-1, NQO-1, or PTEN, respectively, and the ensuing secondary antibodies against IgG arising from distinct species.

### 2.13. Coimmunoprecipitation Analyses

Different plasmids (Keap1-Flag, Keap1α-Flag, Keap1β-Flag and Nrf2-V5) were transfected into HepG2 cells (4 × 10^5^) that had been allowed for 24-h growth on 6-well plates. After 8 h of transfection, the cells were recovered for 24 h in a fresh medium. The experimental cells were lysed in the RIPA buffer (C1053, PPLYGEN) containing both protease and phosphatase inhibitors (Solarbio, P6730). The total lysates were centrifuged, and the resulting supernatants were subjected to coimmunoprecipitation (CO-IP) with antibodies against V5 or Flag (Invitrogen), before being pulled down by the BeyoMag™ Protein A+G beads (Beyotime [24], P2108-1ML). The eluted proteins were separated by SDS–PAGE. The protein-blotted membranes were probed with V5 or Flag antibodies and horseradish peroxidase-conjugated secondary antibodies to IgG (ZSGB-BIO, ZB-2305, 0.1ML), prior to being visualized by enhanced chemiluminescence reagents (Thermo, iBright 750[25]).

### 2.14. Immunofluorescence and Confocal Microscopy

After alcohol-soaked glass coverslips (1 cm^2^) were placed flat in 6-well plates, COS-1 cells (2.5 × 10^5^) were allowed for being grown on the glass-stored plates for 24 h, and then subjected to transfection with each of indicated plasmids expressing Keap1-V5, Keap1α-V5 and Keap1β-V5, respectively. After transfection for 8 h and then recovery for 24 h, the cells were fixed with a paraformaldehyde fixative (Servicebio, Wuhan) for 30 min at room temperature, and then permeabilized with 0.2% Triton X-100 for 20 min. The fixed cells were incubated with 1% BSA for 60 min, and immunoblotted with each of the primary antibodies overnight, and the secondary Alexa Fluor–conjugated secondary antibody (anti-mouse or anti-rabbit antibody, ZSGB-BIO) for 4 h. The glass-mounted cells were then rinsed with PBS three times, before being subjected to DNA-staining with DAPI (Beyotime, C1005). The immunofluorescent mages were achieved by confocal microscopy with an IN Cell Analyzer Zeiss LSM900 cellular imaging system[26].

### 2.15. Transcriptome Sequencing Analysis

Total RNAs, that had been extracted from experimental cells using an RNAsimple total RNA Kit, were subjected to transcriptome sequencing analysis on an Illumina HiSeq 2000 sequencing system (Illumina, San Diego, CA) in the Beijing Genomics Institute (BGI, Shenzhen, China). From the transcriptome data, all those differentially expressed genes (DEGs) were identified by standard fold changes of ≥2 or ≤0.5 with an FDR (false discovery rate) of ≤0.001, using the Poisson distribution model method (PossionDis), and further analyzed in-depth as described previously [22,27].

### 2.16. Subcutaneous Tumour Xenografts in Nude Mice

Two subcutaneous tumour xenograft experiments were conducted in nude mice (all relevant animal experiments complied with the regulations of the Animal Ethics Committee in this study with mice being kept under the conditions of an SPF experimental animal room). In the first experiments, mouse xenograft models were made by subcutaneous heterotransplantation of wild-type *HepG2* (*Keap1^+/+^*), *Keap1^-/-^*, or *Keap1β*(*Keap1^Δ1-31^*) cell lines into nude mice as described previously [28]. In the second experiments, similar mouse xenograft models were also established by subcutaneously heterotransplantating *Keap1^-/-^, Keap1-Restored* or *Keap1α-Restored* cell lines into nude mice. These transplantated cell lines (1.0 ×10^7^, growing in the exponential phase) were each suspended in 0.1 ml of PBS and then inoculated subcutaneously into male nude mice (BALB/cnu/nu, 6 weeks, 15 g, 6 mice per group, from HFK Bioscience, Beijing, China) at a single point in the back area. The sizes of the tumours were subsequently measured every other day. The tumour-bearing mice were respectively sacrificed on the 24th or 30th day in these two distinct experiments, and meantime their transplanted tumours were excised surgically. The tumour sizes were then calculated by a standard formula: V = a (length) × b^2^ (width^2^)/2.

### 2.17. Immunohistochemistry with HE Staining

The above subcutaneous tumours excised from xenograft nude mice were fixed with a paraformaldehyde fixative (Servicebio, Wuhan) for 24 h. The tumour tissues were subjected to immunohistochemical staining with specific antibodies against Keap1 or PTEN, respectively, in addition to routine histopathological examination (HE) by haematoxylin-eosin (HE) staining. The reagents and antibodies used herein were provided by Servicebio.

### 2.18. Statistical Analysis

All statistical significances in cell proliferation, migration, cycle, apoptosis, invasion and gene expression were determined by Student’s *t*-test (for comparisons between two groups) or two-way ANOVA (for comparisons among multiple groups), respectively. The relevant data are obtained from at least three independent experiments and shown as fold changes (mean ± SEM or ± SD, relative to indicated controls), with significant differences being calculated by the value of *p* < 0.05 or 0.01.

## 3. Results

### 3.1. Four model cell lines with stable expression of Keap1^-/-^, Keap1β(Keap1^Δ1-31^)), Keap1-Restored, and Keap1α-Restored were established

The *Keap1^-/-^* cell line was constructed using CRISPR/Cas9 technology, and identified by real-time qPCR and Western blotting to confirm true with no expression of *Keap1* (Figure 1A), and also further validated by sequencing its genomic DNA showing that 10 bases being chopped from alleles of *Keap1* to result in a change in its open reading frame (Figure S1A, B). In the selection process, another cell lines only expressing a smaller band of Keap1 was obtained and designated as *Keap1β*(*Keap1^Δ1-31^*) herein (Figure 1B). Further sequencing revealed an extra base insertion prior to the 23rd base position in the CDS region of *Keap1*, leading to its open-reading frame shift (Figure S1C, D).

To verify this, a full-length sequence plasmid of pLVX-mcmv-ZsGreen-puro pKeap1 was made. Through alignment with the NCBI database, it was found that the second N-terminal translation starting codon ATG of Keap1 is positioned at the 32^nd^ aa residue. It is inferable that the 32^nd^ aa translation initiation signal is critical for Keap1β expression. Thus, the 32^nd^ aa-codon (i.e., the second ATG) was deleted to create pLVX-mcmv-ZsGreen-puro pKeap1α. Through lentivirus infection, two cell lines (*Keap1-Restored* and *Keap1α-Restored*) were constructed on the base of *Keap1^-/-^* cells and confirmed by real-time qPCR and western blotting (Figure 1C). The results demonstrate that the N-terminal 32^nd^ residue position of Keap1 determines the expression of Keap1β. Therefore, differential expression of Keap1 and its isoforms α and β was caused by distinct compilation of its constitutive amino acids. Of note, two key positions are at the 1^st^ and 32^nd^ methionine residues (Figure 1D). Upon deletion of the 32^nd^ residue from Keap1, only Keap1α was forcedly expressed, whereas its N-terminal aa 1-31 were truncated, only Keap1β was allowedly expressed.

**Figure 1.**
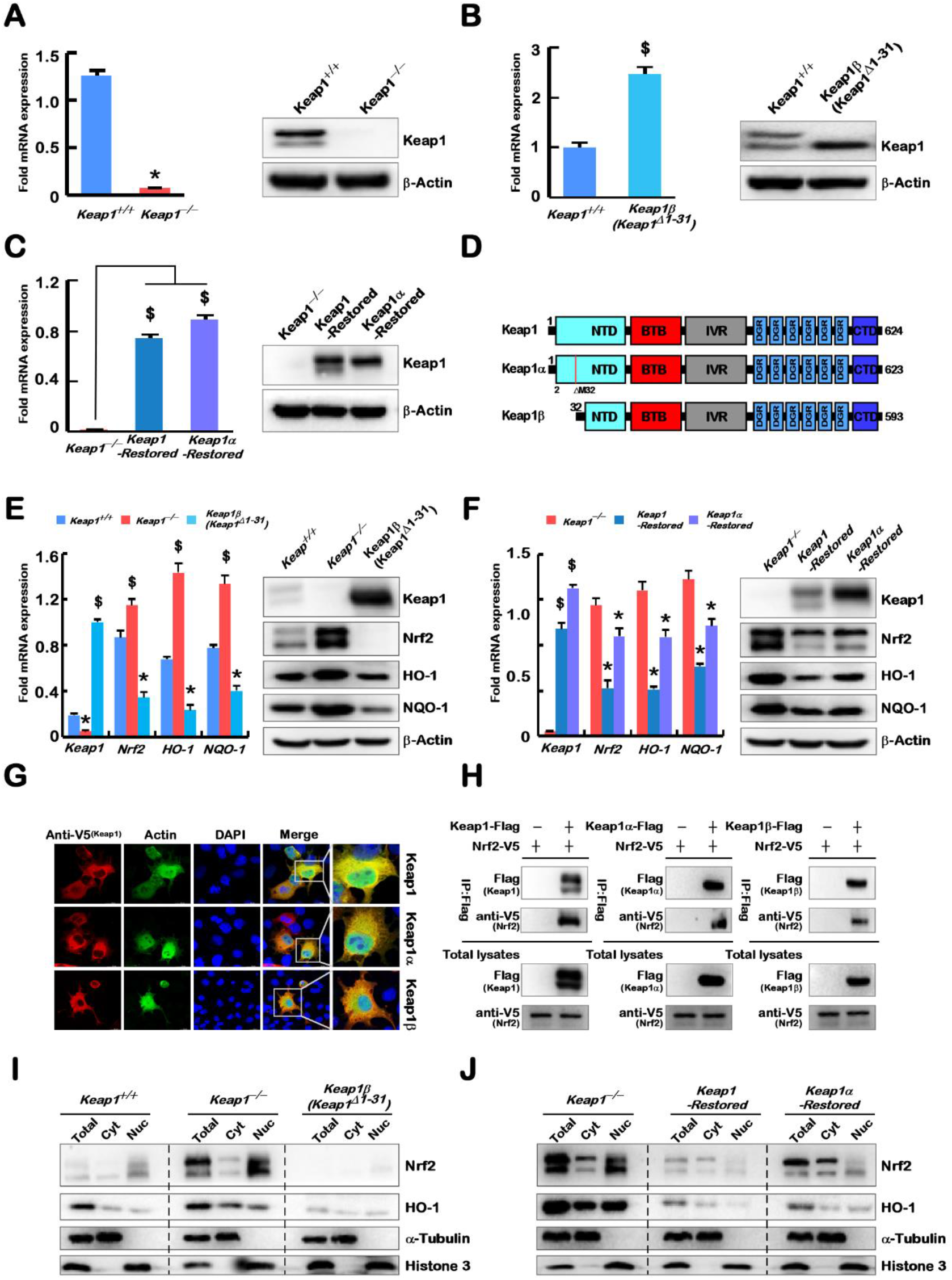
Construction of model cell lines with differential expression of Keap1 and its isoform α and β. **(A-C)** Identification of *Keap1^-/-^* **(A)**, *Keap1β*(*Keap1^Δ1-31^*) **(B)**, *Keap1-Restored* **(C)** and *Keap1α-Restored* **(C)** cell lines by real-time qPCR and Western blotting. **(D)** Schematic representation of Keap1 and its isoform with distinct domains. When compared with Keap1, Keap1α lacks the 2nd translation starting codon (ATG, encoding the 32^nd^ amino acid), whilst Keap1β lacks the first N-terminal 31 amino acids. (E,F) The mRNA and protein expression levels of *Keap1, Nrf2, HO-1* and *NQO-1* were determined in *Keap1^+/+^, Keap1^-/-^* and *Keap1β*(*Keap1^Δ1-31^*) cell lines **(E),** and also examined in *Keap1^-/-^, Keap1-Restored* and *Keap1α-Restored* cell lines (F). **(G)** Subcellular localization of Keap1 and its isoforms α and β in COS-1 cells that had been transfected with Keap1-V5, Keap1α-V5 or Keap1β-V5 plasmids, respectively. Then immunocytochemistry was performed by incubating the cells with primary antibody and fluorescent secondary antibody, before being stained with DAPI. A flag-tagged primary antibody against endogenous Actin was used as a control. The fluorescence intensity of indicated proteins was measured and observed by an immunofluorescence microscope. The images showed a representative of three independent experiments. Scale bars = 20 μm. **(H)** Interaction of Nrf2 with Keap1 and its isoforms α and β was verified by the co-immunoprecipitation (CO-IP) analysis of HepG2 cells that had been co-transfected with Nrf2 and each expression construct for Keap1, Keap1α or Keap1β, respectively. After transfection for 8 h and recovery for 24 h, the total proteins was collected and subjected to CO-IP assays. The resulting immunoprecipitates, along with total cell lysates, were analyzed by immunoblotting with the indicated antibodies. **(I, J)** Subcellular fractionation of Nrf2 together with Keap1 and its isoforms α and β. The cytoplasmic and nuclear fractions were isolated from five distinct cell lines and visualized by Western blotting with antibodies against Nrf2, HO-1 and α-tubulin and histone3. Statistical significant increases ($) or decreases (*) were indicated by *p* ≤ 0.01 (n ≥3, or = 3×3).

By comparison with *Keap1^+/+^*, the expression levels of *Nrf2*, and its target genes *HO-1* and *NQO-1* were increased in *Keap1^-/-^* cells, but decreased in *Keap1β*(*Keap1^Δ1-31^*) cells (Figure 1E). Further comparison of *Keap1^-/-^* with *Keap1-Restored* and *Keap1α-Restored* cells revealed decreases in the expression of both *Nrf2*, *HO-1* and *NQO-1* levels (Figure 1F), and obvious decreases were detected in *Keap1-Restored* cells. Different expression constructs for Keap1-V5, Keap1α-V5 and Keap1β-V5 were transfected into COS-1 cells, before being subjected to immunofluorescence experiments. The results showed that Keap1, Keap1α and Keap1β were all located primarily in the cytoplasm (Figure 1G), whilst the cytoskeleton actin (as a reference control) was present in the nucleus and cytoplasm. Reciprocal immunoprecipitation assays revealed that Nrf2 enabled for interaction with Keap1, Keap1α and Keap1β (Figure 1H). Through nucleocytoplasmic fractionation experiments, it was found that in the absence of Keap1, Nrf2 and HO-1 were upregulated in the cytoplasm, whereas overexpression of Keap1β resulted in their significant downregulation (Figure 1I). Further examinations also unraveled that upon overexpression of Keap1 (*Keap1α-Restored* cells) and Keap1α (*Keap1α-Restored* cells), Nrf2 and HO-1 were significantly downregulated in both the nuclear and cytoplasmic compartments (Figure 1J). Overall, both Keap1α and Keap1β are able to bind and regulate Nrf2 and its downstream genes, though to different degrees.

### 3.2. Differential expression profiles of genes regulated by Keap1^-/-^, Keap1-Restored, Keap1α-Restored and Keap1β (Keap1^Δ1-31^) were defined

Transcriptome sequencing was employed to identify differential expression profiles of genes regulated by *Keap1^-/-^, Keap1-Restored, Keap1α-Restored* and *Keap1β*(*Keap1^Δ1-31^*) in the established model cell lines. From all detectable genes, those upregulated or downregulated with fold changes of ≥2 or ≤0.5, respectively, plus a false discovery rate (FDR) ≤0.001 (Figure 2A), were redefined as differentially expressed genes (DEGs) by comparison with the control data of *Keap1^+/+^* cells under the same conditions. As a result, 1147 DEGs were detected in *Keap1^-/-^* cells, including 756 upregulated genes and 391 downregulated genes (Figure 2A and Table S1). Notably, 887 DEGs were identified in *Keap1-Restored* cells, of which 640 were upregulated and 247 were downregulated (Table S2). In contrast, *Keap1α-Restored* cells also gave rise to 640 upregulated and 247 downregulated genes (Table S3), whereas *Keap1β*(*Keap1^Δ1-31^*) cells led to a significant decreased number of its upregulated genes (i.e., 235), as accompanied by the downregulation of 157 DEGs (Table S4). Together, these data indicate that their DEGs were significantly reduced when only a single isoform α or β of Keap1 were expressed. The common genes between every two cell groups and every specific gene were also selected (Figure 1B), and these comparison groups had distinct differences. Of note, those genes regulated by Keap1, as well as its isoforms α and β, were more likely to be upregulated than downregulated (Figure 1B and Table S3).

The biological process terms and pathways mediated by Keap1 were classified, based on the KEGG and QuickGO databases. Those DEGs were subjected to further analysis of the KEGG pathway and GO enrichment, screening for the top 10 functions for comparison (Figure S2). Such functional annotation analysis implied that DEGs regulated by Keap1 (*cf*. *Keap1^-/-^* with *Keap1-Restored* cell lines) were predominantly involved in cancer, signal transduction, cellular community, infectious disease, signalling molecules and interaction, immune system process, metabolic process, biological adhesion, developmental process, cellular process, biological regulation and cellular process. Similarly, Keap1α-regulated DEGs (in *Keap1α-Restored* cells) were responsible for cancer, endocrine and metabolic disease, immune disease, immune system, substance dependence, cell growth and death, cardiovascular disease, infectious disease, signal transduction, immune system process, biological regulation, cellular process, developmental process, biological regulation, cellular process and biological regulation. By contrast, Keap1β-regulated DEGs (in *Keap1β*(*Keap1^Δ1-31^*) cells) were responsible for endocrine and metabolic disease, cardiovascular disease, cancer, signal transduction, signalling molecules and interaction, infectious disease, developmental process, cell proliferation, biological adhesion, developmental process, biological regulation, positive regulation of biological process, cellular process and biological regulation. From these results, it was inferable that they had highly related functions. As exemplified in Figure S2, the PI3K-AKT pathway was found multiple times in the KEGG pathway analysis.

The crossovers of distinct profiling DEGs regulated by *Keap1^-/-^*, *Keap1-Restored*, *Keap1α-Restored* and *Keap1β*(*Keap1^Δ1-31^*) were displayed in a Venn diagram (Figure 2C). Their specific DEGs were numbered as follows: 297 in *Keap1^-/-^* cells, 68 in *Keap1-Restored* cells, 143 in *Keap1α-Restored* cells, and 166 in *Keap1β*(*Keap1^Δ1-31^*) cells. The top 10 KEGG pathways and GO enrichment functions of these specific DEGs were, in detail, analyzed in another article from our group. Approximately 137 DEGs were screened to be shared among the four cells, as shown in the Venn diagram (Figure 2C and Table S5). The differential expression levels of these common DEGs (137) were presented by differential clustering heatmaps (Figure 2D). These genes were also subjected to biological process analysis. Their biological processes mainly focus on cellular processes, metabolic processes, biological regulation, positive regulation of biological processes and responses to stimuli. Amongst all the shared DEGs, a majority of them showed similar expression trends (both up- and downregulated) in all four cell lines, but only 36 genes were manifested in inconsistent trends, (Figure 2D). Thereby, the functions of Keap1, Keap1α and Keap1β overlap greatly, albeit there are still some differences between them.

**Figure 2.**
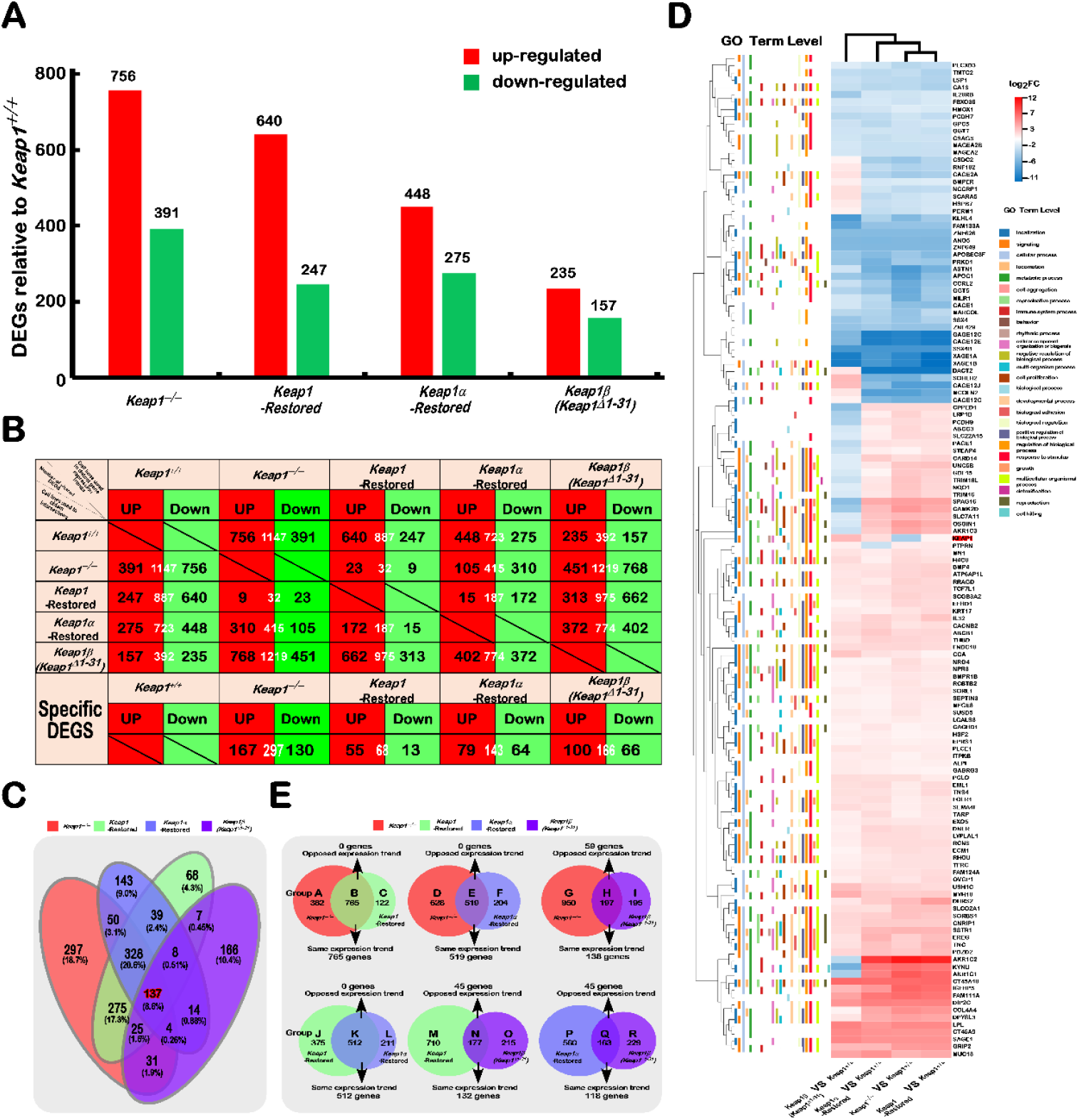
Statistical analysis of the data obtained from transcriptome sequencing. **(A)** The differentially expressed genes (DEGs) of distinct cell lines were analyzed by transcriptome sequencing, and the increase or decrease in DEGs was presented in the form of a histogram The DEGs were selected according to the following criteria: fold changes of ≥2 or≤0.5 and FDR ≤0.001 (as compared to those obtained from control cells). **(B)** The specific DEGs in each cell line and their common DEGs between every two cell lines were also counted as indicated in the chart, and the number of increased and decreased DEGs in each group is shown separately in black font, and the total is shown in white. In addition, the change trends of DEGs in each group were indicated in red or green, which represent upregulated or downregulated in the cells (*in the first row*), respectively. **(C)** The common or unique DEGs among sequenced samples were exhibited in the Venn chart. **(D)** The heatmap with hierarchical clustering of 137 DEGs was shared in all four cells lines, when compared to the data of *Keap1^+/+^* cell lines. **(E)** Venn diagrams represent those common and differentially expressed genes between every two cells.

### 3.3. Functional annotation of specific or common DEGs in Keap1^-/-^, Keap1-Restored, Keap1α-Restored and Keap1β(Keap1^Δ1-31^) cells

To gain an in-depth understanding of the similarities and differences in biological functions between *Keap1^-/-^, Keap1-Restored, Keap1α-Restored* and *Keap1β*(*Keap1^Δ1-31^*), their common and different regulatory DEGs were identified in different combinations of each pair of groups, as indicated by Groups A to R (Figure 2E and Table S6). The similar or opposite trends in those common DEGs of every two cell groups are shown. For example, 765 DEGs between *Keap1^-/-^* and *Keap1-Restored* cells had the similar expression trend to that as shown in Group B. In Group E, there was only an expression trend among the 595 DEGs shared between *Keap1^-/-^* and *Keap1α-Restored* cells. Two distinct expression trends coexisted in Group H, in which 59 DEGs expressed different trends and 138 DEGs had the same expression trend in *Keap1^-/-^* and *Keap1β*(*Keap1^Δ1-31^*) cells. In Group K, 512 DEGs between *Keap1-Restored* and *Keap1α-Restored* cells had the same expression trend. In contrast, largely identical expression trends were observed in 132 of the common 177 DEGs coregulated by *Keap1 -Restored* and *Keap1β*(*Keap1^Δ1-31^*) and in 118 of the other common 163 DEGs shared by *Keap1α-Restored* and *Keap1β*(*Keap1^Δ1-31^*) cells in Groups N and Q.

To determine the differences between *Keap1^-/-^, Keap1-Restored, Keap1α-Restored* and *Keap1β* (*Keap1^Δ1-31^*) cells, the top 10 KEGG pathways and GO enrichment with DEGs were exhibited in scatterplots and histograms. The mRNA expression levels of selected genes were determined by real-time qPCR, and the putative functions of these target genes were mapped (Figure S3-S8 and Table S6). All DEGs regulated by Keap1 alone or both were divided into three groups, A, B and C, and then functionally annotated with the above-described methods (Figure S3A and Table S6). In Group A, 382 DEGs regulated in *Keap1^-/-^* cells were generally involved in signal transduction, cancer, infectious disease, transport and catabolism, metabolic processes and cellular processes. In the intersected Group B, 765 DEGs co-regulated in both *Keap1^-/-^* and *Keap1-Restored* cell lines were also associated with cardiovascular disease, signal transduction, biological adhesion, metabolic process and cellular process. In Group C, 122 DEGs regulated in *Keap1-Restored* cells were significantly enriched in the cellular community, endocrine system, cancer, signal transduction, immune system process, localization and developmental process. Moreover, the expression of *AKR1C3, ALP1, GDP15, HPD, HSPB1, ID3, PGD, PHGDH* and *SESN2* was validated by real-time qPCR (Figure S3B). In addition, putative functions of such target genes were mapped, as indicated by the histogram (Figure S3B, *on the bottom*).

The common and distinct targeting DEGs in *Keap1^-/-^* and/or *Keap1α-Restored* cell lines were assigned to three groups. In Group D, 628 DEGs were identified as targets of *Keap1^-/-^* cells and were enriched with distinct functions in signal transduction, cellular community, infectious disease, biological regulation, metabolic process and biological regulation (Figure S4A and Table S6). Their 519 shared DEGs in Group E were functionally responsible for cancer, cardiovascular disease, signal transduction, immune system processes, biological regulation and metabolic processes. In Group F, 204 DEGs were regulated preferentially in *Keap1α-Restored* cells and functionally associated with substance dependence, cancer, signal transduction, biological regulation, cellular processes and developmental processes. Amongst those DEGs, the expression of *EPHX-1, HISTIHIC, HLA-A, IARS, SAT1, SESN2, SMAD3* and *TFRC* was further validated by real-time qPCR (Figure S4B). The putative functions relative to these examined genes are illustrated (in Figure S4B, *on the bottom*).

Those DEGs regulated in *Keap1^-/-^* and/or *Keap1β*(*Keap1^Δ1-31^*) cell lines in Groups G to I were functionally annotated by DAVID, and histograms and scatterplots were herein constructed (Figure S5A and Table S6). Further comprehensive analysis of the 10 top significant KEGG pathways and the enriched biological process terms showed that 950 DEGs of Group G, from *Keap1^-/-^* cells, were associated with distinct functions in signal transduction, infectious disease, cancer, biological regulation, the cellular process and developmental process, etc. In Group H, across *Keap1^-/-^* and *Keap1β*(*Keap1^Δ1-31^*), 197 co-regulated DEGs were identified that were involved in cancer, signaling molecules and interaction, signal transduction, cellular process, metabolic process and developmental process. In Group I, only 195 DEGs was identified by transcriptional regulation by *Keap1β* (*Keap1^Δ1-31^*). Their putative functions were associated with cancer, endocrine and metabolic disease, signal transduction, biological regulation, biological adhesion, and positive regulation of biological processes. The expression of *CD44, FGFBP1, HISTIH2BK, ID1, IL20RB, MYC, SESN2, SGK1, TALDO1* and *TFG-β1* were validated by real-time qPCR (Figure S5B). The putative functions relative to these examined genes are shown (in Figure S5B, *on the bottom*).

The common and distinct target DEGs in *Keap1-Restored* and/or *Keap1α-Restored* cell lines were assigned to three groups. In Group J, 375 DEGs were identified as targets of *Keap1-Restored* cells and were enriched with distinct functions in cardiovascular disease, signal transduction, cell motility, cancer, immune system process, biological adhesion, metabolic process and cellular process (Figure S6A and Table S6). The 512 DEGs shared in Group K were functionally responsible for cancer, endocrine and metabolic diseases, signaling molecules and interactions, signal transduction, immune system processes, cellular processes and immune system processes. In Group L, 211 DEGs were regulated preferentially by *Keap1α-Restored* cells and functionally associated with cancer, substance dependence, cell growth and death, signal transduction, cellular processes, biological regulation and developmental processes. Further real-time qPCR validated the expression of CLDN1, *HLA-C, OPN3, SESN2, SMAD3, SMAD4, SOAT1, SQSTM1, TGF-βl* and *TNPO1* (Figures S6B). The putative functions relative to these examined genes are shown (in Figure S6B, *on the bottom*).

All DEGs regulated in *Keap1-Restored* and/or *Keap1β Keap1^Δ1-31^*) cell lines were divided into Groups M to O and then functionally annotated with the above-described methods (Figure S7A and Table S6). In Group M, 710 DEGs regulated in *Keap1-Restored* cells were generally involved in signal transduction, immune system, cancer, immune system process, biological adhesion, metabolic process and developmental process. In the intersected Group N, 177 DEGs coregulated by *Keap1-Restored* and *Keap1β*(*Keap1^Δ1-31^*) were associated with development and regeneration, cancer, signal transduction, signaling molecules and interaction, signal transduction, biological adhesion, developmental process, the cellular process and biological regulation. In Group O, 215 DEGs regulated by *Keap1β*(*Keap1^Δ1-31^*) were significantly enriched in endocrine and metabolic disease, cancer, infectious disease, signal transduction, biological regulation, developmental process, positive regulation of biological process and cell proliferation. Real-time qPCR results validated that expression of *AKR1B1, AKR1C1, BLVR8, BMP2, FTL, G6PD, HISTIHIC, HLA-C, PI3, PSAT1, SMAD2, SQSTM1, TFRC* and *TGF-β1* (Figures S7B). In addition, putative functions of such target genes were mapped, as indicated by the histogram (Figure S7B, *on the bottom*).

The DEGs regulated by *Keap1α-Restored* and/or *Keap1β*(*Keap1^Δ1-31^*) in Groups P, Q and R were functionally annotated by DAVID, and histograms and scatterplots were also constructed (Figure S8A and Table S6). Comprehensive analysis of the 10 top significantly KEGG pathways and the enriched biological process terms revealed that 560 DEGs of Group P, in *Keap1α-Restored* cells, were associated with distinct functions in infectious disease, substance dependence, immune disease, signal transduction, immune system process, biological regulation, the cellular process and developmental process. In Group Q, *Keap1α-Restored* and *Keap1β*(*Keap1^Δ3-31^*) coregulated 163 DEGs that were involved in cancer, development and regeneration, signaling molecules and interaction, endocrine and metabolic disease, developmental process, biological adhesion and cellular process. In Group R, only 229 DEGs were identified by transcriptional regulation in *Keap1β*(*Keap1^Δ1-31^*) cells. Their putative functions were associated with endocrine and metabolic diseases, signal transduction, signaling molecules and interactions, cancer, biological regulation, developmental processes, cell proliferation and positive regulation of biological processes. The results of real-time qPCR validated the expression of *CTSB*, *GARS, GDF15, HIST1H4H, LAMB3, S100A16, SMAD2, TFRC, TGF-β1* and *TWIST2* (Figure S8B). The putative functions relative to these examined genes are shown (in Figure S8B, *on the bottom*).

Collectively, those genes regulated by *Keap1^-/-^, Keap1-Restored, Keap1α-Restored* and *Keap1β*(*Keap1^Δ1-31^*) were different, but notably the number of genes regulated by Keap1α was greater than that regulated by Keap1β. A comprehensive analysis of the four cell lines found their similar functions in KEGG pathways and GO enrichment as follows: cancer, signal transduction, signaling molecules and interaction, biological regulation, the cellular process and developmental process. There were also different KEGG pathways and GO enrichment functions: cellular community, endocrine system, immune system, infectious disease, localization, immune system process and response to the stimulus. Of note, the PI3K-mTOR pathways monitored by both Keap1 isoforms were found in the KEGG pathway analysis.

### 3.4. Keap1 knockout mutant was a more potent player than Keap1β(Keap1^Δ1-31^) at preventing tumour xenografts in nude mice

To investigate the effect of deletion of all Keap1 or Keap1α alone on tumour growth, *Keap1^-/-^* and *Keap1β* (*Keap1^Δ1-31^*), along with *Keap1^+/+^* (wild-type, *WT*), cell lines were heterotransplanted into three groups of immunodeficient nude mice at their subcutaneous loci as indicated. After tumour formation, the sizes of growing tumours were measured at one-day intervals within the ensuing 25 days, before the tumour-bearing mice were sacrificed. The resulting data of tumour in sizes and weights were calculated and graphed. The resulting images showed that the carcinomas derived from *Keap1^+/+^* cells were estimated to be ~7.0 times in tumour volume and weight larger than those of *Keap1^-/-^* cells and 3.0 times than that of *Keap1β*(*Keap1^Δ1-31^*) cells (Figure 3A-C). The fact that the carcinomas derived from *Keap1β* were larger than carcinomas derived from *Keap1^-/-^* and litter than carcinomas derived from *Keap1^+/+^* demonstrates that *Keap1* and partially *Keap1β*(*Keap1^Δ1-31^*) can promote the occurrence and development of tumours.

Then, xenograft tumours were subjected to histopathological examination by routine haematoxylin-eosin (HE) staining, followed by immunohistochemical staining with antibodies against Keap1 and PTEN (Figure S9). Through HE staining experiments, it was observed that some cell necrosis and even liquefaction occurred in the tumours formed by *Keap1^+/+^* and *Keap1β*(*Keap1^Δ1-31^*) cell lines, among which *Keap1^+/+^* cells had more necrosis, whereas there was almost no similar pathology in tumours formed by *Keap1^-/-^* cells (Figure S9A). In the tumour-bearing tissue formed by *Keap1^-/-^* cells, the overall colouration of the cells was darker, and there were a large number of cancer grooves in the formed tumours to wrap the tumour cells.

When compared with *Keap1^+/+^* cells, the expression of Keap1 was markedly decreased in *Keap1^-/-^-derived* tumour tissues, but markedly increased in *Keap1β*(*Keap1^Δ1-31^)-derived* tumour tissues (Figure S9B). Further immunohistochemistry of the tumour-repressing marker PTEN revealed significant enhancements of PTEN in *Keap1^-/-^* and *Keap1β*(*Keap1^Δ1-31^*) derived xenograft tissues (Figure S9C). Furtherly, a series of comparative experiments revealed that cell proliferation levels were also significantly suppressed by either *Keap1^-/-^* or *Keap1β* (*Keap1^Δ1-31^*) mutants, as compared with the *Keap1^+/+^* control (Figure 3D).

**Figure 3.**
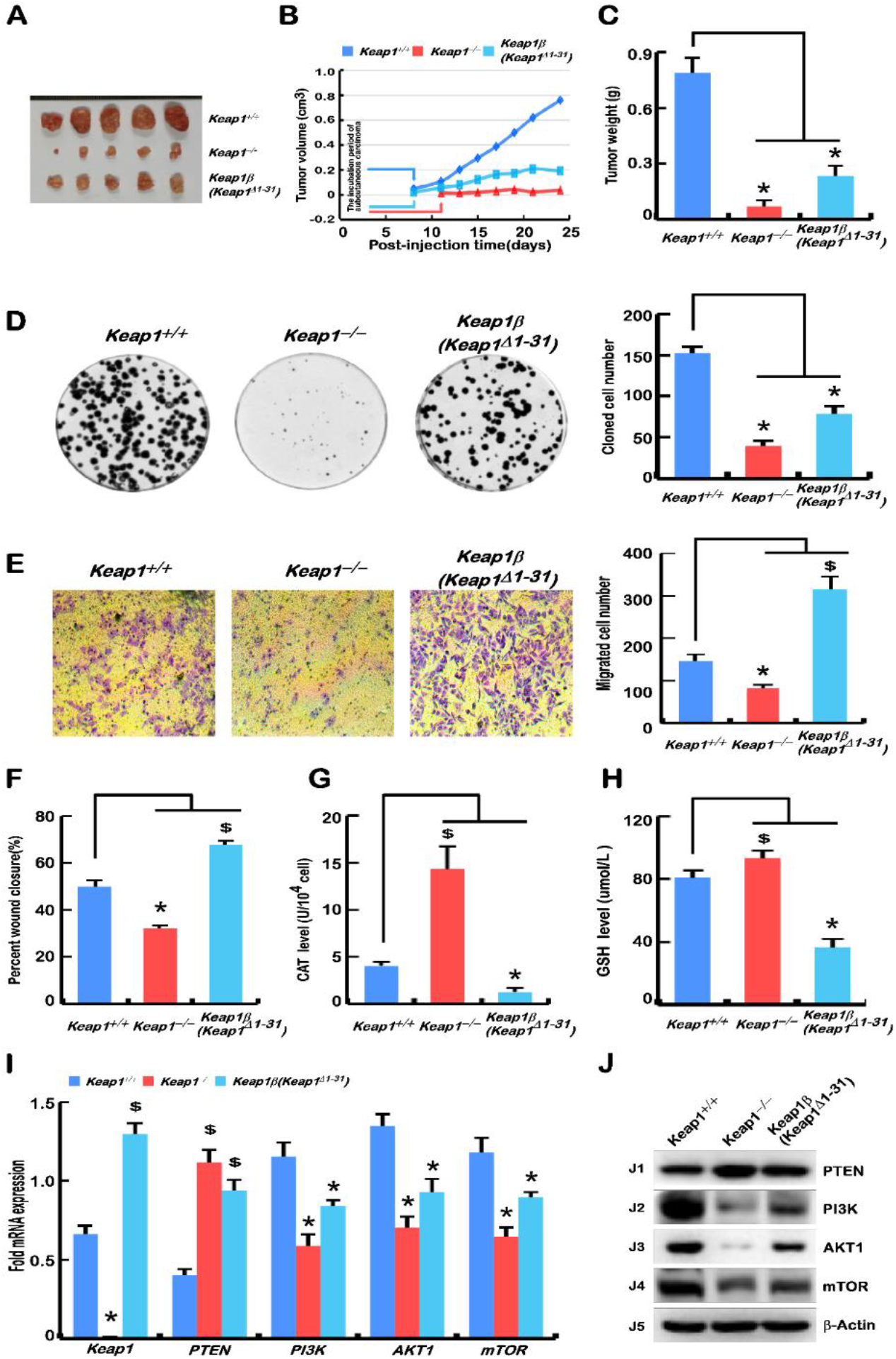
Knockout of Keap1 is a more potent player than *Keap1β*(*Keap1^Δ1-31^*) at preventing tumour xenografts in nude mice. **(A)** *Keap1^+/+^, Keap1^-/-^* and *Keap1β*(*Keap1^Δ1-31^*) cell lines were respectively injected subcutaneously in each group of 5 nude mice, and their tumours were fetched out after growing for 25 days. **(B)** Tumour growth curves of *Keap1^+/+^, Keap1^-/-^* and *Keap1β*(*Keap1^Δ1-31^)-derived* carcinomas in subcutaneous tissues of nude mice within 25 days. **(C)** These tumour weights were measured after removal from the subcutaneous tumour-bearing loci of nude mice. **(D)** Cell proliferation was calculated by counting the colony-forming units. Equal amounts of *Keap1^+/+^, Keap1^-/-^* and *Keap1β*(*Keap1^Δ1-31^*) cell lines were cultured in each capsule, and 10 days later, their corresponding clones was quantified. **(E)** The number of those cells migrated across the Transwell membrane in *Keap1^+/+^, Keap1^-/-^* and *Keap1β*(*Keap1^Δ1-31^*) cell lines were calculated, after the polycarbonate film stained with crystal violet was photographed on an inverted microscope. **(F)** Wound healing was assessed according to the scratch width, as also shown in Figure S10A and S10B. The cell mobility of each of indicated lines was calculated at 24 h, 48 h and 72 h. **(G)** The enzymatic activity of CAT in *Keap1^+/+^, Keap1^-/-^* and *Keap1β*(*Keap1^Δ1-31^*) cell lines was determined according to the manufacturer’s instruction. **(H)** The intracellular GSH levels of *Keap1^+/+^, Keap1^-/-^* and *Keap1β*(*Keap1^Δ1-31^*) cell lines were measured according to the relevant instruction. (**I)** The mRNA expression levels of *Keap1, PTEN, PI3K, AKT1* and *mTOR* in *Keap1^+/+^, Keap1^-/-^* and *Keap1β*(*Keap1^Δ1-31^*) cell lines were determined by real-time qPCR. **(J)** The protein abundances of Keap1, PTEN, PI3K, AKT1 and mTOR expressed in *Keap1^+/+^, Keap1^-/-^* and *Keap1β*(*Keap1^Δ1-31^*) cell line were visualized by Western blotting. Statistical significant increases ($) or decreases (*) were indicated by *p* ≤ 0.01 (n ≥3, or = 3×3).

The cell cycle results showed that *Keap1^-/-^* cells underwent a cycle arrest at G1 phase, which is contributable to tumour reduction, but no significant changes in *Keap1β* cell cycle were observed (Figure S10C). Meanwhile, apoptosis of *Keap1^-/-^* and *Keap1β* cell lines was increased (Figure S10D). Further experiments showed that the invasion and migration of *Keap1^-/-^* cells were inhibited, while those of *Keap1β*(*Keap1^Δ1-31^*) cells were promoted (Figures 3E, 3F, S10A, S10B). The relevant data of CAT activity along with intracellular GSH levels were showed that they were increased in *Keap1^-/-^* cells, but decreased in *Keap1β*(*Keap1^Δ1-31^*) cells, as compared with the control of *Keap1^+/+^* cells (Figure 3G, 3H).

Based on the xenograft tumour results, real-time qPCR and Western blotting were performed to reexamine the tumour repressor PTEN to mTOR signaling pathways. The results obtained from *Keap1^-/-^* and *Keap1β*(*Keap1^Δ1-31^*) cells unraveled that PTEN was obviously upregulated, whilst PI3K, AKT1 and mTOR were all downregulated (Figures 3I, 3J), showing the similar trends to those of *in vivo* growing tumours (Figure 3,A-C). However, their changes (increased or decreased) were more pronounced in the *Keap1*-expressing cell lines. From these, it is inferable that *Keap1 and Keap1β* may function as more potent tumour promoters, albeit both isoforms were endowed with an intrinsic capability to inhibit hyper-active Nrf2 at preventing tumour development and malgrowth. Such opposing effects of *Keap1 and Keap1β* depends on their monitored activity of the PTEN signaling to the PI3K, AKT1 and mTOR pathway, besides Nrf2-regulated ARE gene networks.

### 3.5. Malgrowth of Keap1^-/-^-derived hepatoma cells was significantly suppressed by the restoration of Keap1α, and the Keap1-Restored promoted their growth

To further explore the effects of Keap1 and Keap1α on tumour growth, *Keap1-Restored* and *Keap1α-Restored* cells, alongside *Keap1^-/-^* (WT) cells, were respectively heterotransplanted into three groups, each including 5 immunodeficient nude mice, at their subcutaneous loci as indicated. After tumour formation, the sizes of growing tumours were measured at one-day intervals within the ensuing 30 days before these tumour-bearing mice were sacrificed. The tumour sizes and weights were calculated as shown graphically. The results obtained were as follows: the human carcinoma derived from *Keap1^-/-^* tumours were estimated to be <u>only 3 times smaller</u> than those derived from *Keap1-Restored* cells, but also 4.0 times larger than those derived from *Keap1α-Restored* cells (Figure 4A-C). These data further demonstrate the potential role of Keap1 in promoting tumour growth; however, it is surprising that Keap1α has an opposite effect on tumour growth as done by Keap1, which can promote tumour progression.

Then, xenograft tumours were subjected to histopathological examination by routine haematoxylin-eosin staining, followed by immunohistochemical staining with antibodies against Keap1 and PTEN (Figure S11). The staining results showed that some cell necrosis and liquefaction occurred in the tumour tissures formed by *Keap1^-/-^, Keap1-Restored* and *Keap1α-Restored* cells, respectively (Figure S11A). When compared with tumours derived from *Keap1^-/-^* cells, in the tumour-bearing tissues formed by *Keap1-Restored* and *Keap1α-Restored* cells, the overall colouration of these cells was darker, which seemed to be more basophilic, and there were a large number of cancer grooves in the tumour formed by *Keap1α-Restored* cells, which inhibited the tumour growth (Figure S11A). Further immunohistochemical examinations revealed significant enhancements of Keap1 in the *Keap1-Restored* and *Keap1α-Restored* derived xenograft tissues (Figure S11B), and the expression of PTEN was markedly decreased in the *Keap1-Restored* cell-derived tumour tissue, but also strikingly increased in the *Keap1α-Restored* cell-derived tumour tissue (Figure S11C).

Further experiments showed that *Keap1^-/-^* cell proliferation, invasion and migration were all significantly promoted by restoration of *Keap1*, while *Keap1α-Restored* cells were significantly suppressed (Figure 4D-F, S12A, S12B). In *Keap1-Restored* cell, the G1 phase of the cell cycle was shortened (Figure S12C). Meanwhile, the restoration of *Keap1* reduced apoptosis, whereas the restoration of *Keap1α* significantly increased apoptosis (Figure S12D). The data from measuring CAT and GSH levels showed that both were decreased in *Keap1-Restored* and *Keap1α-Restored* cells as compared with those obtained from *Keap1^-/-^* cells (Figures 4G, 4H).

Such obvious phenotypes of xenograft tumourwere further were validated by real-time qPCR and Western blotting analyses of the expression of the tumour-specific marker PTEN-mTOR pathway. The results from the *Keap1-Restored* cells showed that *PTEN* was downregulated, whilst *PI3K*, *AKT1* and *mTOR* were upregulated, but a completely opposite trend was found in *Keap1α-Restored* cells (Figures 4I, 4J).

As a consequence, the restoration of Keap1α enabled a significant tumour-preventing effect on subcutaneous human carcinoma xenografts in nude mice, but the recovery of Keap1 promoted the tumour effect. Collectively, it is inferable that Keap1 acts as a potent tumour promoter, and its major isoform Keap1α can prevent tumour development and malgrowth.

**Figure 4.**
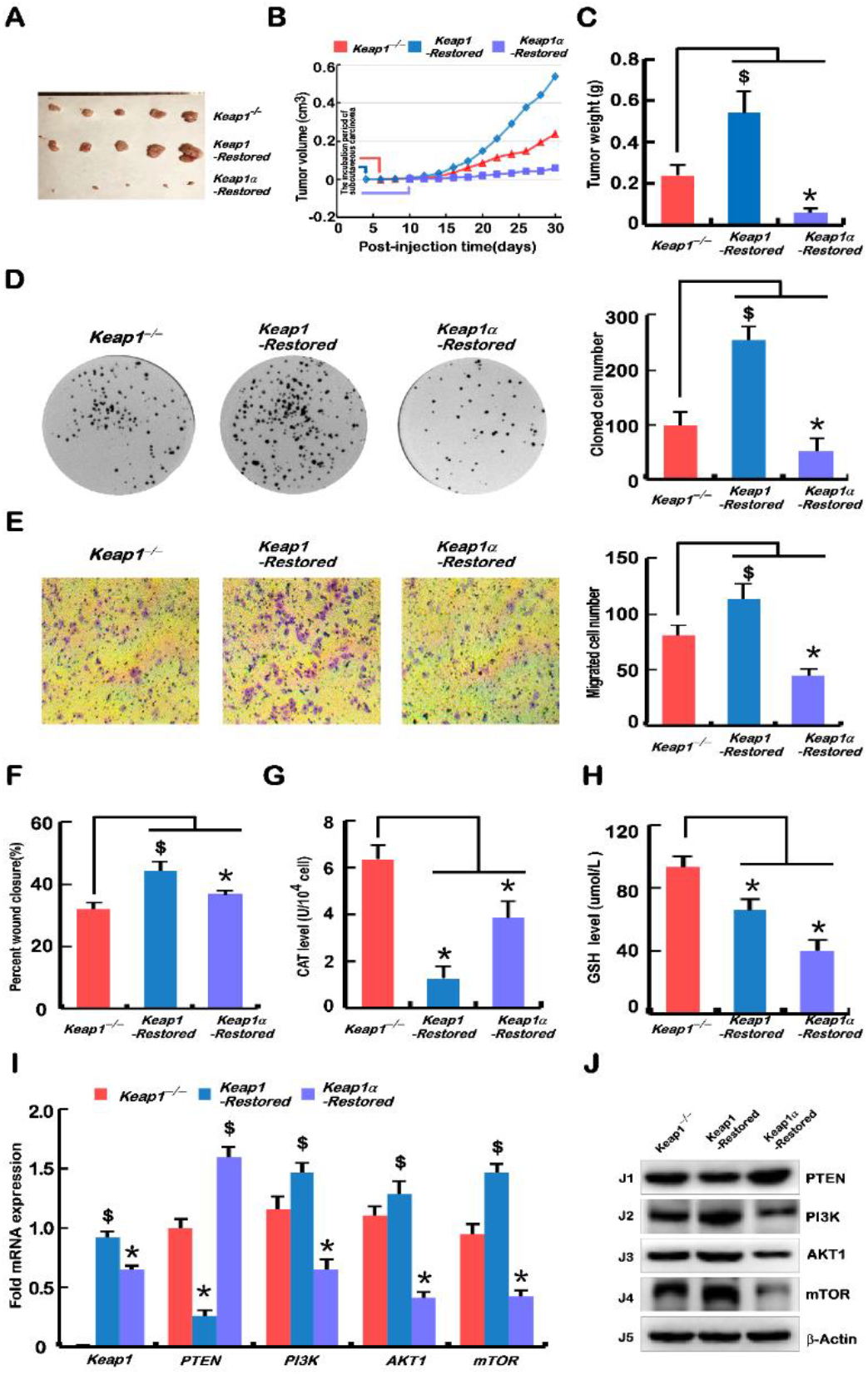
Malgrowth of *Keap1^-/-^*-derived hepatoma cells were significantly suppressed by restoration of Keap1α, but the *Keap1-Restored* can promote their growth. **(A)** Subcutaneous tumour formation in nude mice. *Keap1^-/-^, Keap1-Restored* and *Keap1α-Restored* cell lines were injected subcutaneously in grouped nude mice, and 30 days later the tumours were fetched out. **(B)** Tumour growth curves of *Keap1^-/-^, Keap1-Restored* and *Keap1α-Restored* cells in subcutaneous tissues of nude mice within 30 days. **(C)** The tumours were excised from the subcutaneous carcinomas loci in nude mice and then weighed. **(D)** Cell proliferation was detected by the colony-forming unit assays. The equal amounts of *Keap1^-/-^, Keap1-Restored* and *Keap1α-Restored* cell lines were cultured in a capsule, and 10 days later the corresponding number of clones in each dish was recorded. **(E)** Transwell migration was assayed. The polycarbonate film stained with crystal violet was photographed on an inverted microscope. The number of cells migrated across the membrane in *Keap1^-/-^, Keap1-Restored* and *Keap1α-Restored* cell lines were calculated. **(F)** Wound healing was assayed according to the scratch width in figure S12A and S12B, before the mobility of each cell was calculated at 24 h, 48 h and72 h. **(G)** Measure of the CAT activity in *Keap1^-/-^*, *Keap1-Restored* and *Keap1α-Restored* cell lines according to the kit instruction. **(H)** Determination of the intracellular GSH levels in *Keap1^-/-^*, *Keap1-Restored* and *Keap1α-Restored* cell lines according to the instruction. (**I)** Real-time qPCR analysis of the mRNA expression levels of *Keap1, PTEN, PI3K, AKT1* and *mTOR* in *Keap1^-/-^, Keap1-Restored* and *Keap1α-Restored* cell lines. **(J)** Western blotting examination of the protein abundances of Keap1, PTEN, PI3K, AKT1 and mTOR in *Keap1^-/-^, Keap1-Restored* and *Keap1α-Restored* cell lines. Statistical significant increases ($) or decreases (*) were indicated by *p* ≤ 0.01 (n ≥3, or = 3×3).

### 3.6. Deficiency of Keap1 and its subtypes (α and β) results in dysregulation of genes controlling cell behaviour

The aforementioned results demonstrated that loss of Keap1 and its subtypes (α and β) led to marked phenotypic changes in cell behaviours, such as tumorigenesis of the derived carcinoma xenografts, and growth, invasion, migration, transformation, cycle and apoptosis from *Keap1^+/+^, Keap1^-/-^, Keap1-Restored, Keap1α-Restored* and *Keap1β*(*Keap1^Δ1-31^) cell lines*. The subset of genes that control cell processes and behaviours was dysregulated in several cell lines, and further identified by real-time qPCR and Western blotting.

In the epithelial-mesenchymal transition (EMT) experiments, *Keap1^+/+^* cells were used as a control. As anticipated in Figure 5A and 5B, the results revealed the upregulation of key genes encoding the epithelial marker proteins *CTNNA1* (*α-Catenin*), *CTNNB1* (*β-Catenin*) and *CDH1* in *Keap1^-/-^* cells. Their upregulation was accompanied by the downregulation of critical genes encoding the mesenchymal marker proteins *CDH2, FN1* and vimentin. Then, the expression levels of zinc finger transcription factors *SNAI1* and *SNAI2* were reduced. Notably, *SNAI1* and *SNAI2* have been shown to act as pivotal mediators of EMT by regulating the expression of its target genes *CDH1* and *CDH2* through binding the E-boxes within their promoter regions, particularly in metastatic hepatocellular carcinoma [23,29,30]. Moreover, loss of Keap1 also led to downregulated expression (at transcriptional and translational levels) of genes encoding membrane-type *MMP17* and matrix metallopeptidase 9 (MMP9) in *Keap1^-/-^* cells. However, *Keap1β*(*Keap1^Δ1-31^*) cells showed completely opposite results to those obtained from *Keap1^-/-^* cells.

When compared with *Keap1^-/-^* cells, real-time qPCR was employed to detect typical genes related to EMT in the *Keap1-Restored* and *Keap1α-Restored* cells. Herein, it was found that the *CTNNA1*, *CTNNB1* and *CDH1* genes were downregulated, while the remaining genes were enhanced, in *Keap1-Restored* cells (Figure 5C). However, the *Keap1α-Restored* cells demonstrated the opposite results of the *Keap1-Restored* cells. In these cells (*Keap1^-/-^, Keap1-Restored* and *Keap1α-Restored cells*), Western blotting examination also showed the similar trends to that measured by real-time qPCR (Figure 5D).

### 3.7. Knockout of Keap1 and its subtypes (α and β) results in dysregulation of genes controlling the cell cycle and apoptosis

To gain insights into the above-mentioned cell cycle and apoptosis changes from the mechanism, the key genes involved in these processes were examined by real-time qPCR. As anticipated, the expression levels of *P53*, *P21* and *CDK2* were significantly downregulated in *Keap1^-/-^* and *Keap1β* (*Keap1^Δ1-31^*) cell lines, as compared with equivalent values obtained from *Keap1^+/+^* control cells (Figure 4E). This finding supported the notion that the S phase of *Keap1^-/-^* and *Keap1β*(*Keap1^Δ1-31^*) cell cycles were reduced. But, it is rather intriguing that *Keap1^-/-^* and *Keap1β*(*Keap1^Δ1-31^*) cell lines gave rise to upregulated mRNA expression levels of genes encoding *CDK6* and *Cyclin D1* (both were involved in a functional complex controlling the checkpoint of G1-S transition) in *Keap1^-/-^* and *Keap1β*(*Keap1^Δ1-31^*) cell lies (Figure 5E).

Further examination revealed that proapoptotic genes (e.g., *Caspases 3, 4* and *6*) were increased, whilst anti-apoptotic genes (e.g., *Bcl2* and *Bcl2l1*) were decreased, in *Keap1^-/-^* and *Keap1β*(*Keap1^Δ1-31^*) cell lines when *Keap1^+/+^* cells served as a reference (Figure 5F). To verify the effects of Keap1 on cell cycle progression and apoptosis, *Keap1-Restored* and *Keap1α-Restored* cells were compared to *Keap1^-/-^* cells to determine the expression levels of related genes. Upregulation of the *P53-P21-CDK2* signaling pathway in *Keap1-Restored* and *Keap1α-Restored* lines led to an increase in the S phase of cell cycles, whereas *CDK6* and *CyclinD*1 were both decreased, possibly to retard the G1-S transition (Figure 5G). Those genes (e.g., *Caspases 3, 4* and *6*) that promote apoptosis were downregulated, and genes (e.g., *Bcl2* and *Bcl2l1*) that resist apoptosis were upregulated, in the *Keap1-Restored* cells, but such proapoptotic and antiapoptotic genes showed the opposite trends of their expression levels in *Keap1α-Restored* cells (Figure 5H).

**Figure 5.**
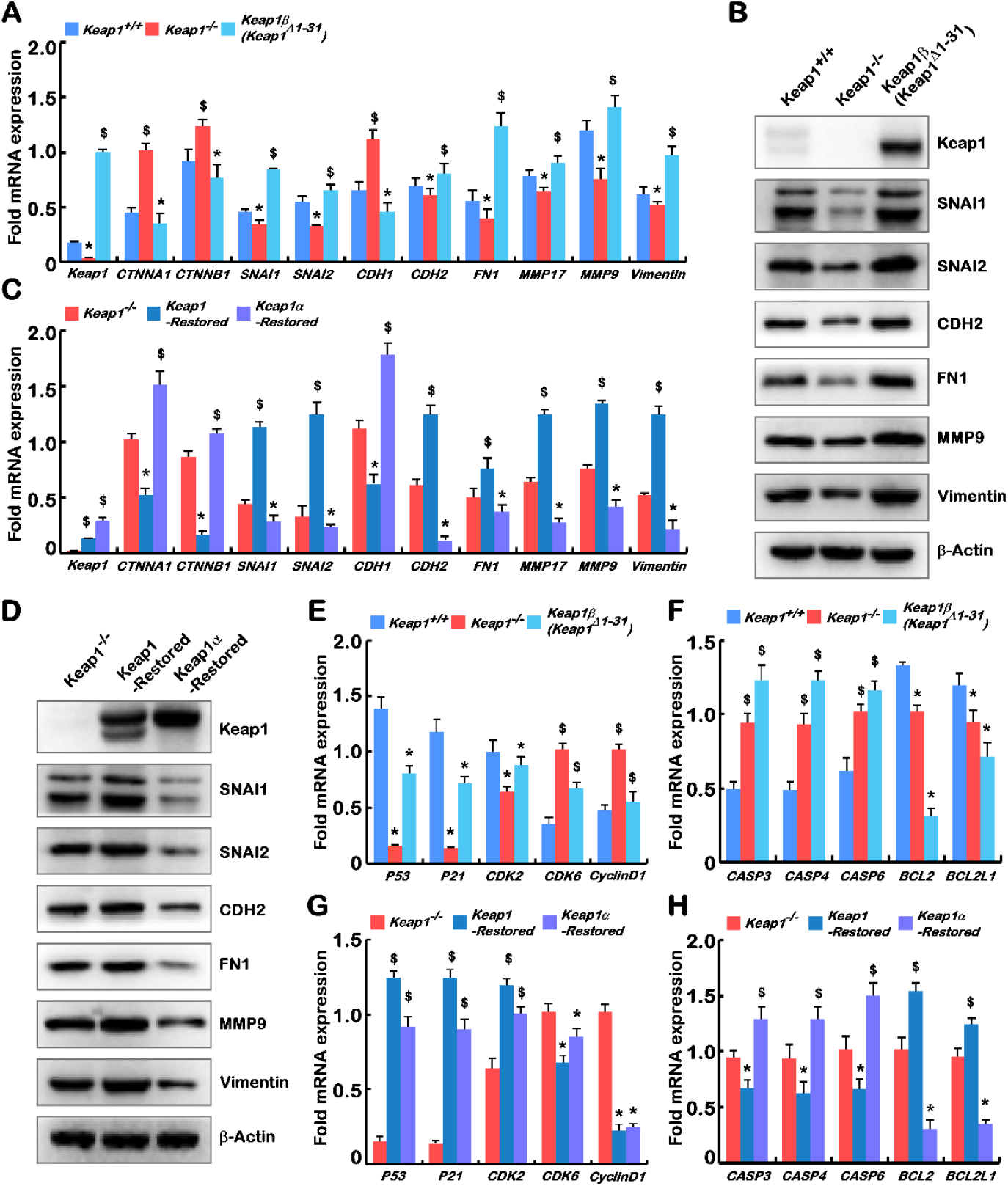
Deficiency of Keap1 and its isoforms α and β lead to gene dysregulation of cell behaviour, cycle and apoptosis. (**A)** Real-time qPCR was used to quantify the mRNA expression levels of *Keap1, CTNNA1, CTNNB1, SNAI1, SNAI2, CDH1, CDH2, FN1, MMP17, MMP9* and *Vimentin* in *Keap1^+/+^, Keap1^-/-^* and *Keap1β*(*Keap1^Δ1-31^*) cell lines. **(B)** Western blotting was employed to determine the protein abundances of Keap1, CTNNA1, CTNNB1, SNAI1, SNAI2, CDH1, CDH2, FN1, MMP17, MMP9 and Vimentin in *Keap1^+/+^, Keap1^-/-^* and *Keap1β*(*Keap1^Δ1-31^*) cell lines. (**C)** The mRNA expression levels of *Keap1, CTNNA1, CTNNB1, SNAI1, SNAI2, CDH1, CDH2, FN1, MMP17, MMP9* and *Vimentin* in *Keap1^-/-^, Keap1-Restored* and *Keap1α-Restored* cell lines were determined by real-time qPCR. **(D)** The protein abundances of Keap1, CTNNA1, CTNNB1, SNAI1, SNAI2, CDH1, CDH2, FN1, MMP17, MMP9 and Vimentin in *Keap1^-/-^, Keap1-Restored* and *Keap1α-Restored* cell lines were visualized by Western blotting with indicated antibodies. (**E)** The mRNA expression levels of genes controlling cell cycle, i.e. *P53, P21, CDK2, CDK6* and *CyclinD1* in *Keap1^+/+^, Keap1^-/-^* and *Keap1β*(*Keap1^Δ1-31^*) cell lines were quantified by real-time qPCR. (**F)** The mRNA levels of genes controlling cell apoptosis, i.e. *CASP3, CASP4, CASP6, BCL2* and *BCL2L1* in *Keap1^+/+^, Keap1^-/-^ and Keap1β*(*Keap1^Δ1-31^*) cell lines were evaluated by real-time qPCR. (**G)** Real-time qPCR was also subjected to determining the mRNA expression levels of *P53*, *P21*, *CDK2, CDK6* and *CyclinD1* in *Keap1^-/-^, Keap1-Restored* and *Keap1α-Restored* cell lines. (**H)** The mRNA expressions of *CASP3, CASP4, CASP6, BCL2* and *BCL2L1* in *KeapT ‘^-^, Keap1-Restored* and *Keap1α-Restored* cell lines were also measured. Statistical significant increases ($) or decreases (*) were indicated by *p* ≤ 0.01 (n ≥3, or = 3×3).

## 4. Discussion

In the present study, we have established four model cell lines allowing for stable expression of *Keap1^-/-^, Keap1β*(*Keap1^Δ1-31^*), *Keap1-Restored* or *Keap1α-Restored*, the latter three of which occur naturally with their respective intact structural domains. Of striking note, the two isoforms of Keap1α and Keap1β were mainly affected by the expression of the N-terminal amino acids 1 and 32, which belong to the NTD domain, but their Ketch/DGR and C-terminal domains are remaining unchanged. Thereby, our experimental evidence further revealing that two isoforms of Keap1 (α and β) could still inhibit the expression of Nrf2 and its target genes HO-1 and NQO-1 in cancer cells (Figure 1E-F). This supports the previously-reported notion that the Ketch/DGR and C-terminal domains of Keap1 directly interacts with the N-terminal Neh2 of Nrf2 and inhibits this CNC-bZIP factor activity [1]. Meanwhile, two isoforms α and β of Keap1 can segregate Nrf2 to be confined in the cytoplasmic subcellular compartments, possibly by its direct interaction with this factor to target the CNC-bZIP protein to the ubiquitin-mediated proteasal degradation pathway, thus inhibiting its expression abundance (Figure 1G-J). In fact, another similar report had also unraveled that overexpression of Keap1a and Keap1b in embryos showed no significant differences in the inhibition of Nrf2[31].

In zebrafish, studies showed that knockout of Keap1a and Keap1b had certain differences in their regulatory genes and pathways, and this were further identified by RNA-sequencing analysis[4]. Here, according to the analysis of transcriptome sequencing data, it was found that the deletion and restoration of Keap1, Keap1α and Keap1β had different regulated genes, among which Keap1 regulated the most and Keap1β the least (Figure 2A, 2B). Venn diagrams and cluster heatmaps were compared, and it was again verified that Keap1 and its isoforms (α and β) had different regulatory functions (Figure 2C-E). The top 10 KEGG pathways and GO enrichments were analyzed in depth, and after pairwise comparison, it was found that their stably-expressing cell lines had similar but different biological processes. Notably, the PI3K-AKT-mTOR signaling pathway had been repeatedly identified in these cell line transcriptome data analysis (KEGG pathways), suggesting that Keap1 and its isoforms (α and β) had a key regulatory role in this pathway (Figure S2-S8).

The topic about the role of Keap1 deletion and subsequent Nrf2 activation is highly debated in the scientific community. Keap1 loss-of-function and/or Nrf2 gain-of-function mutations were predictors of poor clinical outcome in different tumour aetiologies, such as in HCC, and ovarian, lung, endometrial, breast, and head and neck cancers[32,33]. For HCC, it has been reported that mutations in Keap1 or Nrf2 have been detected in approximately 12% of cases[33,34], implying that an active Keap1-Nrf2 pathway could induce or even drive the development of HCC. Conversely, the study of a mouse model constructed with Keap1 knockout in the liver had showed smaller tumours[33]. Of note, our data also showed that *Keap1^-/-^* xenograft mice developed fewer and smaller tumours than *Keap1^+/+^* control mice, with a lower tumour frequency in nude mouse subcutaneous tumour transplantation experiments (Figure 3A-C). Additionally, we observed that tumours recovered by *Keap1α-Restored* cells were smaller than those recovered by *Keap1^-/-^* cells (Figure 4A-C). This suggests that complete intact Keap1 promotes tumour growth, but as Keap1 isoforms, Keap1α inhibits tumour growth, whilst Keap1β promotes tumour growth. The opposite effects of two different subtypes of Keap1 and their differential expression proportions in distinct tissues may be an axiomatic reason why Keap1 plays dual tumour-promoting and tumour-suppressing roles in different tissues. This is also confirmed by further supportive experiments, revealing that deletion of Keap1 or Keap1 isoforms (α or β) inhibited tumour cell colony formation *in vitro* (Figure 3D, 4D).

It is known that more than 90% of cancer deaths are related to tumour cell metastasis[35], and the EMT is generally considered to drive cancer invasion and metastasis during the malignant transformation of liver carcinomas[36,37]. Herein, our evidence has also been provided that deletion of Keap1 inhibited the *in vitro* migration and invasion of cancer cells, whereas Keap1β expression alone showed the opposite results (Figure 3E, 3F, S10A, S10B). Restoring Keap1 expression promoted the *in vitro* migration and invasion of cancer cells, whereas Keap1α expression alone had the opposite effects (Figure 4E, 4F, S12A, S12B). The above experimental results demonstrate that the EMT behaviour of these cell lines was differentially shaped. Thereby, it is reasoned that Keap1 and its isoforms (α or β) affected cell migration and the expression of adhesion junction genes (Figure 5A-D). In our experiments, it was found that Keap1 and its isoforms (α and β) had different effects on these relevant gene expression profiles. Further experimental evidence has also been presented, unveiling that deletion of Keap1 led to activated expression of CAT and increased GSH levels; when only Keap1 isoforms α or β were present, their expression levels were suppressed (Figure 3G, 3H, 4G, 4H). Collectively, the experimental results showed that Keap1 and its isoforms (α and β) had opposite regulatory effects on these activities.

As a matter of fact, the formation of tumours is affected by multiple carcinogenetic pathological factors, among which the PTEN signaling to the PI3K-AKT-mTOR pathway has been studied in more depth. This is due to the fact that PTEN is identified as a typical tumour suppressor that inhibits the expression of the PI3K-AKT-mTOR pathway, and hence regulates cell proliferation, apoptosis, protein synthesis and glucose metabolism, and promotes tumour progression [38]. Overexpression of PI3K in hepatocytes could lead to steatosis and lipid accumulation, accelerating tumour formation[39,40]. The similar experimental evidence was also obtained here by immunohistochemistry (Figure S9C, S11C), real-time PCR (Figure 3I, 4I) and Western blotting (Figure 3J, 4J). More specifically, the PI3K-AKT-mTOR signaling activation could also trigger tumour initiation and cancer cell proliferation and by regulating cell cycle progression [41,42]. A series of comparative experiments by us showed that the S phase of *Keap1^-/-^* cell cycles was the shortest, but the S phase was modestly increased when only expression of *Keap1α-Restored* or *Keap1β*(*Keap1^Δ1-31^*) was present (Figure S10C, S12C). Our evidence further demonstrates that the deletion of Keap1 or the presence of an isoform (α or β) significantly increased the apoptosis rate, and the apoptosis rate was the highest when only its isoform α was present (Figure S10D, S12D). Meanwhile, our results also revealed that Keap1 and its isoforms (α and β) deregulated the expression levels of cell cycle and apoptosis genes (Figure 5E-H). These were basically consistent with the results of *in vitro* experiments measuring the cell cycle and apoptosis by flow cytometry. Taken together, it is inferable that Keap1 and its isoforms (α and β) affect the expression of cell cycle- and apoptosis-related genes regulated possibly by monitoring the PTEN signaling to the PI3K-mTOR pathway.

In summary, we have herein provided a series of compelling *in vitro* and *in vivo* evidence, demonstrating that Keap1 and its subtypes (α and β) have similar but different functions in tumour formation, regulation of the EMT-based invasion and migration, as well as cell cycle progression and apoptosis of metastatic cells. It should be noted that the effects of Keap1 deletion and subsequent activation of Nrf2 are differentially manifested in distinct tumour models, and thus these results are still controversial.

## Author Contributions

Both F.C. and M.X. performed the experiments with help of J.F., R.W., K.L. and S.H., collected all the relevant data, and wrote a draft of this manuscript with most figures and supplemental information. Y.Z. designed and supervised this study, analyzed all the data, helped to prepare all figures with cartoons, rewrote and revised the paper. All coauthors have read and agreed to the published version of the manuscript.

## Acknowledgments

We are greatly thankful to Professor Guiyin Sun (Chongqing University Jiangjin Hospital) for his guidance on the experimental methods in this study. We also thank all other present and past members of Zhang’s laboratory (at Chongqing University, China) for giving critical discussion and invaluable help with this work.

