## supplemental Files for "Different inhibition of Nrf2 by two Keap1 isoforms α and β to shape malignant behaviour of human hepatocellular carcinoma" (from another version of this preprint: version 2, 2022-07-16)

#### Supplemental materials:

#### Figure S1

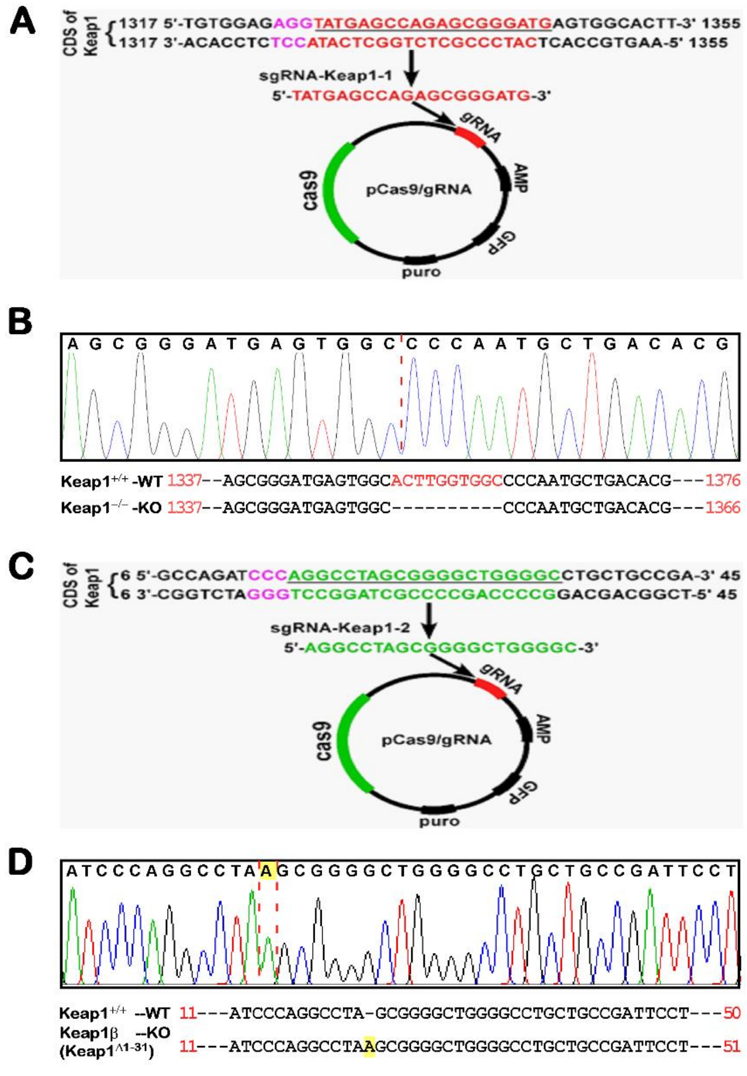


**Figure S1.** Identification of cells. **(A)** Cas9 plasmid used in *Keap1*$^{-/-}$ cells construction. **(B)** Sequence peak diagram of editing site in *Keap1*$^{-/-}$ cells. **(C)** Cas9 plasmid used in *Keap1β(Keap1*$^{Δ1-31}$*)* cells construction. **(D)** Sequence peak diagram of editing site in *Keap1β(Keap1*$^{Δ1-31}$*)* cells.

### Figure S2

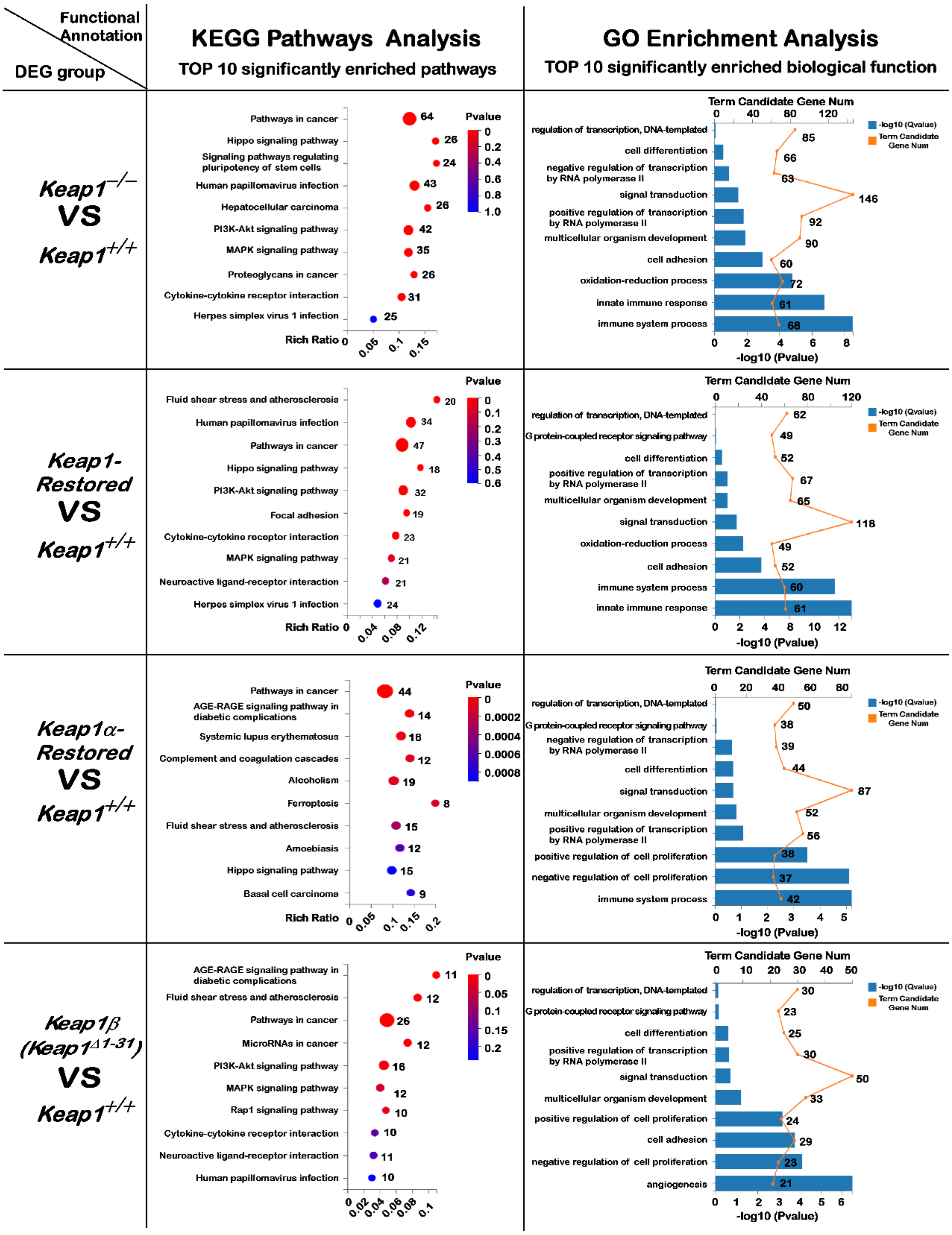


**Figure S2.** Functional annotation of specific or common DEGs in *Keap1*$^{-/-}$, *Keap1-Restored*, *Keap1α-Restored* and *Keap1β(Keap1*$^{\Delta 1\text{-}31}$*)* cells. Through transcriptome data analysis, *Keap1*$^{+/+}$ were compared with the four cells, and the differential genes of each group were screened out. According to the DEGs in each group, KEGG pathway and GO enrichment analysis were performed. The top 10 of significant pathways and biological process terms enriched by DEGs were exhibited in scatterplots and histograms, respectively. Numbers indicate the number of genes.

### Figure S3

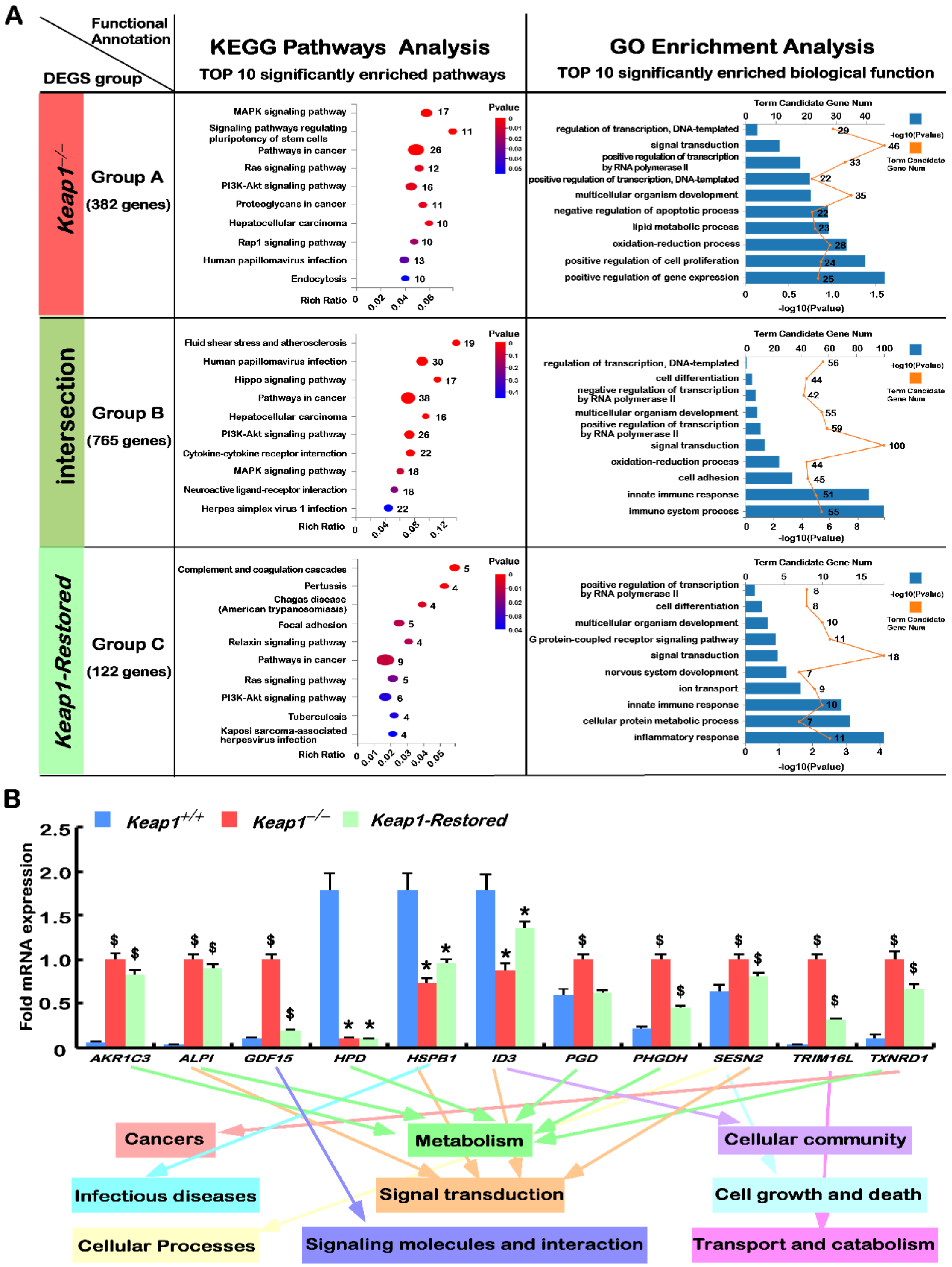


**Figure S3.** Functional annotation of specific or common DEGs in *Keap1*$^{-/-}$ and *Keap1-Restored* cells. **(A)** The top 10 of significant KEGG pathways terms and GO enriched by DEGs in Groups A, B, and C were exhibited in scatterplots and histograms, respectively. Numbers indicate the number of genes. **(B)** Q-PCR was used to detect the mRNA expressions of *AKR1C3, ALP1*, *GDP15*, *HPD*, *HSPB1*, *ID3*, *PGD*, *PHGDH* and *SESN2* in *Keap1*$^{+/+}$, *Keap1*$^{-/-}$ and *Keap1-Restored* cells. Putative functions of such target genes were mapped, as indicated by the histogram. The data are shown as mean±SEM((n=3X3, and "*" "$" indicates p≤0.01).

### Figure S4

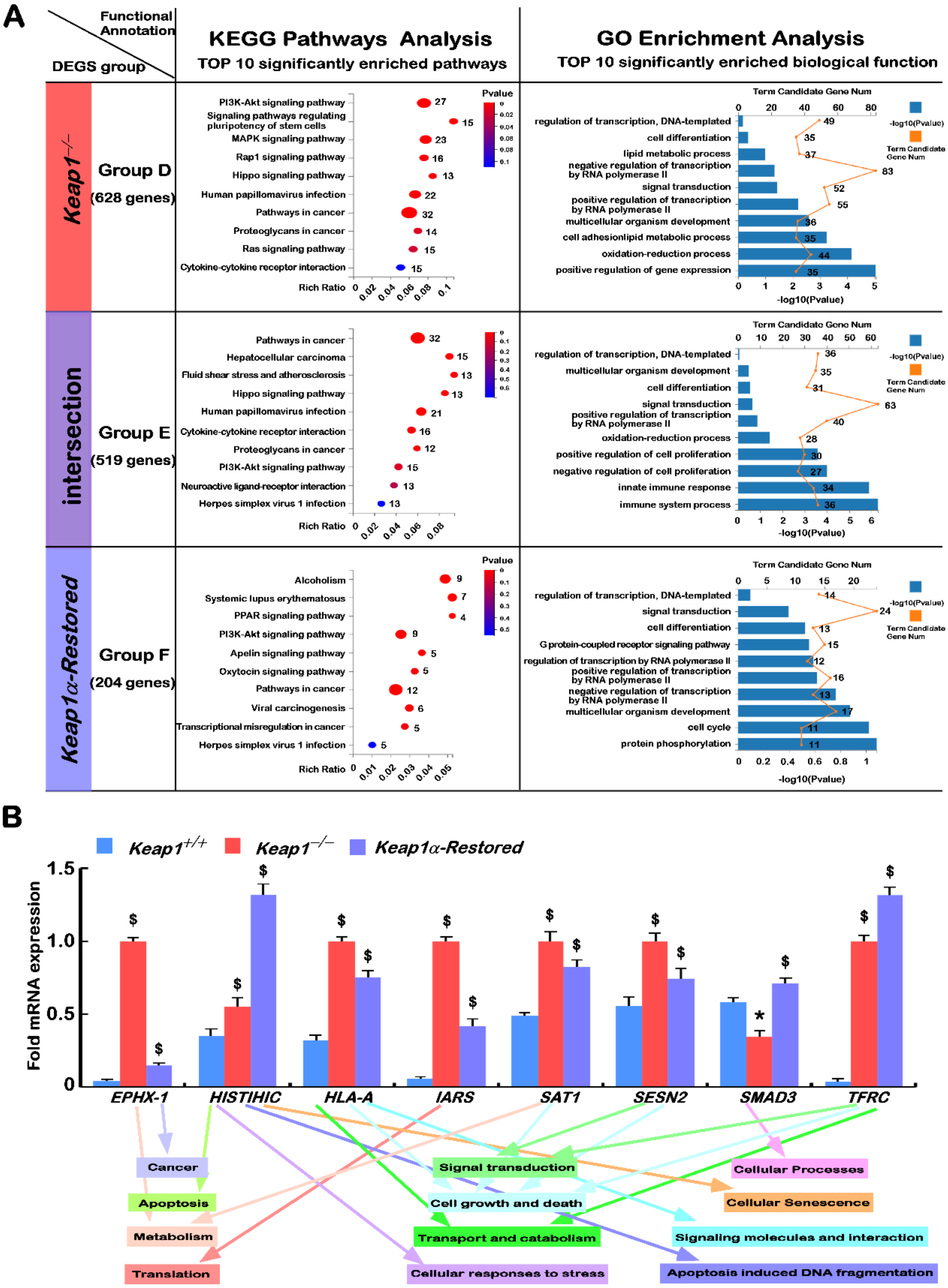


**Figure S4.** Functional annotation of specific or common DEGs in *Keap1*$^{-/-}$ and *Keap1α-Restored* cells. **(A)** The top 10 of significant KEGG pathways terms and GO enriched by DEGs in Groups D, E, and F were exhibited in scatterplots and histograms, respectively. Numbers indicate the number of genes. **(B)** Q-PCR was used to detect the mRNA expressions of *EPHX-1*, *HISTIHIC*, *HLA-A*, *IARS*, *SAT1*, *SESN2*, *SMAD3* and *TFRC* in *Keap1*$^{+/+}$, *Keap1*$^{-/-}$ and *Keap1α-Restored* cells. Putative functions of such target genes were mapped, as indicated by the histogram. The data are shown as mean±SEM(n=3X3, and "*" "$" indicates p≤0.01).

### Figure S5

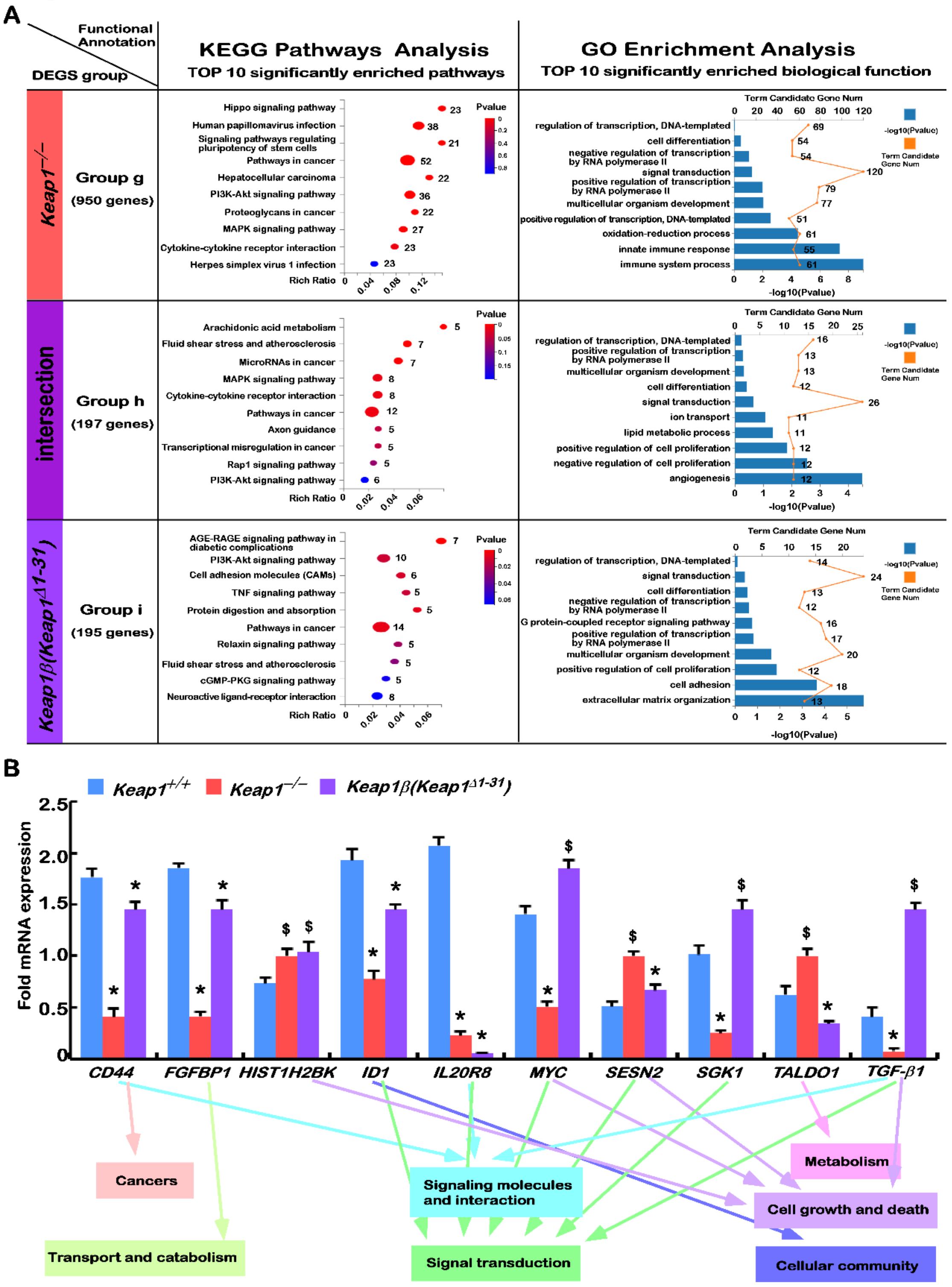


**Figure S5.** Functional annotation of specific or common DEGs in *Keap1*$^{-/-}$ and *Keap1β(Keap1*$^{\Delta 1\text{-}31}$*)* cells. **(A)** The top 10 of significant KEGG pathways terms and GO enriched by DEGs in Groups G, H, and I were exhibited in scatterplots and histograms, respectively. Numbers indicate the number of genes. **(B)** Q-PCR was used to detect the mRNA expressions of *CD44,FGFBP1,HISTIH2BK,ID1,IL20RB,MYC, SESN2, SGK1,TALDO1* and *TFG-β1* in *Keap1*$^{+/+}$, *Keap1*$^{-/-}$ and *Keap1β(Keap1*$^{\Delta 1\text{-}31}$*)* cells. Putative functions of such target genes were mapped, as indicated by the histogram. The data are shown as mean±SEM(n=3X3, and "*" "$" indicates p≤0.01).

### Figure S6

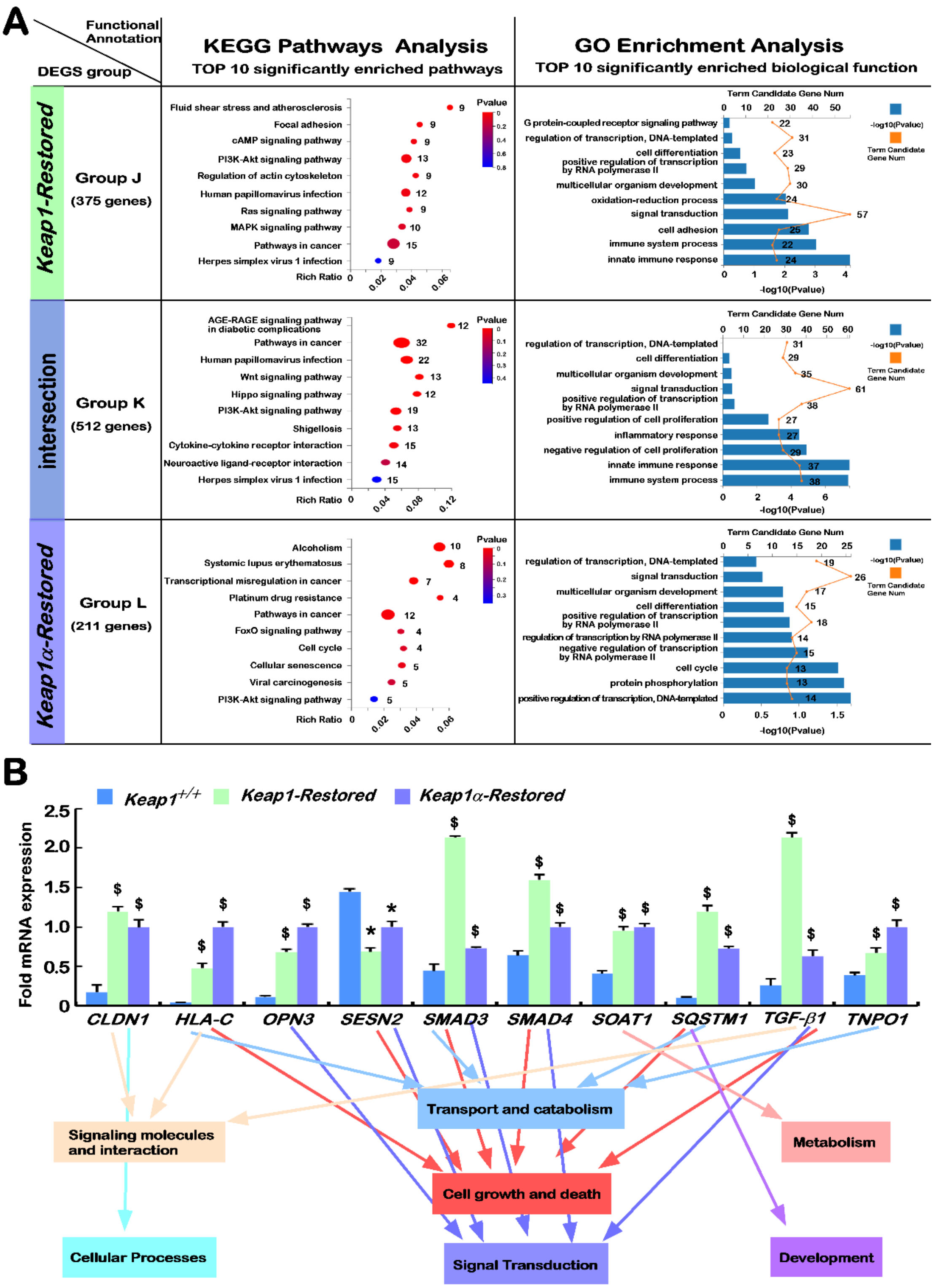


**Figure S6.** Functional annotation of specific or common DEGs in *Keap1-Restored* and *Keap1α-Restored* cells. **(A)** The top 10 of significant KEGG pathways terms and GO enriched by DEGs in Groups J, K, and L were exhibited in scatterplots and histograms, respectively. Numbers indicate the number of genes. **(B)** Q-PCR was used to detect the mRNA expressions of *CLDN1,HLA-C,OPN3,SESN2, SMAD3, SMAD4, SOAT1,SQSTM1,TGF-β1* and *TNPO1* in *Keap1*$^{+/+}$, *Keap1-Restored* and *Keap1α-Restored* cells. Putative functions of such target genes were mapped, as indicated by the histogram. The data are shown as mean±SEM(n=3X3, and "*" "$" indicates p≤0.01).

### Figure S7

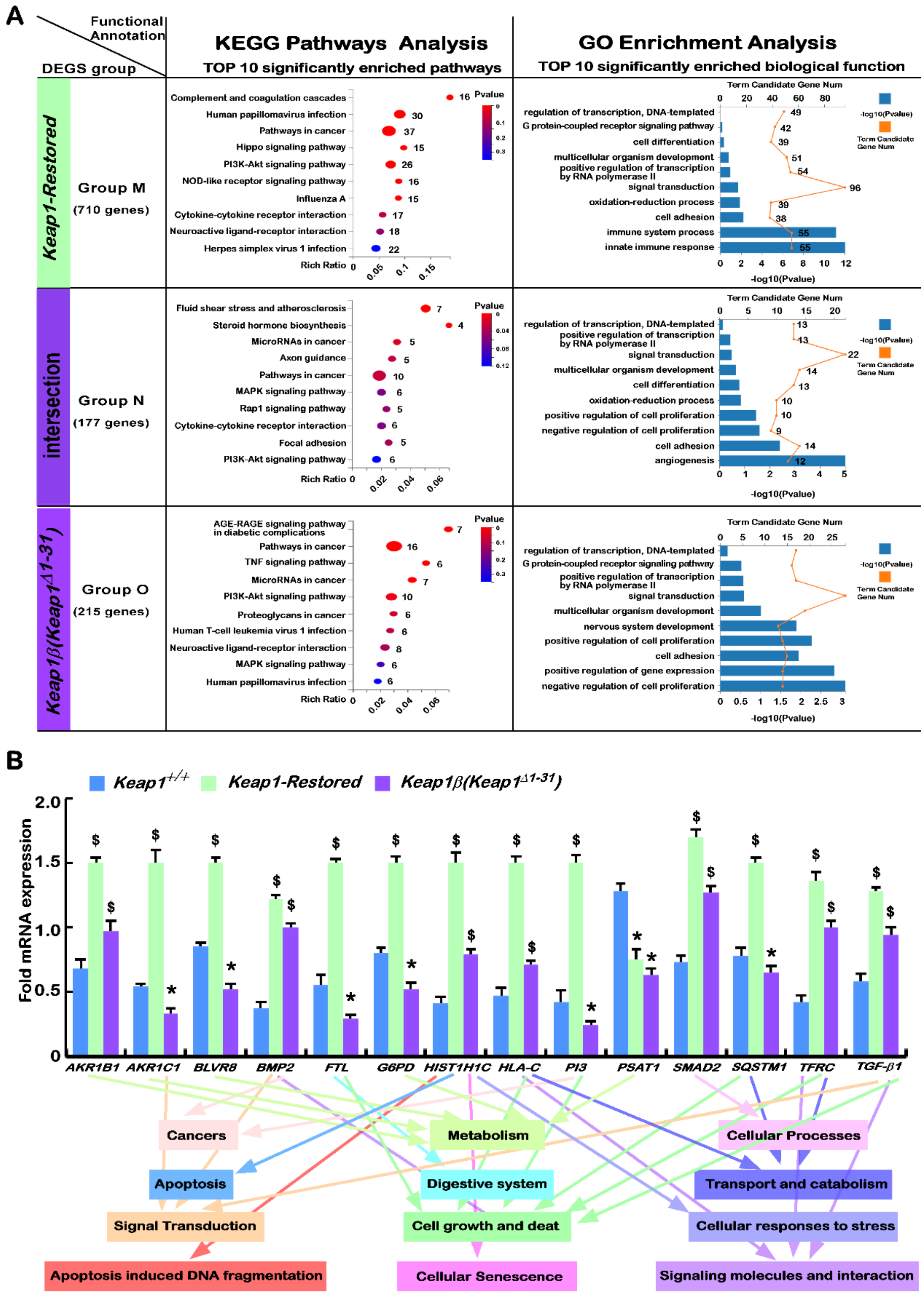


**Figure S7.** Functional annotation of specific or common DEGs in *Keap1-Restored* and *Keap1β(Keap1$^{Δ1-31}$)* cells. **(A)** The top 10 of significant KEGG pathways terms and GO enriched by DEGs in Groups M, N, and O were exhibited in scatterplots and histograms, respectively. Numbers indicate the number of genes. **(B)** Q-PCR was used to detect the mRNA expressions of *AKR1B1,AKR1C1,BLVR8, BMP2,FTL,G6PD,HISTIHIC,HLA-C,PI3,PSAT1,SMAD2,SQSTM1,TFRC* and *TGF-β1* in *Keap1$^{+/+}$*, *Keap1-Restored* and *Keap1β(Keap1$^{Δ1-31}$)* cells. Putative functions of such target genes were mapped, as indicated by the histogram. The data are shown as mean±SEM(n=3X3, and "*" "$" indicates p≤0.01).

### Figure S8

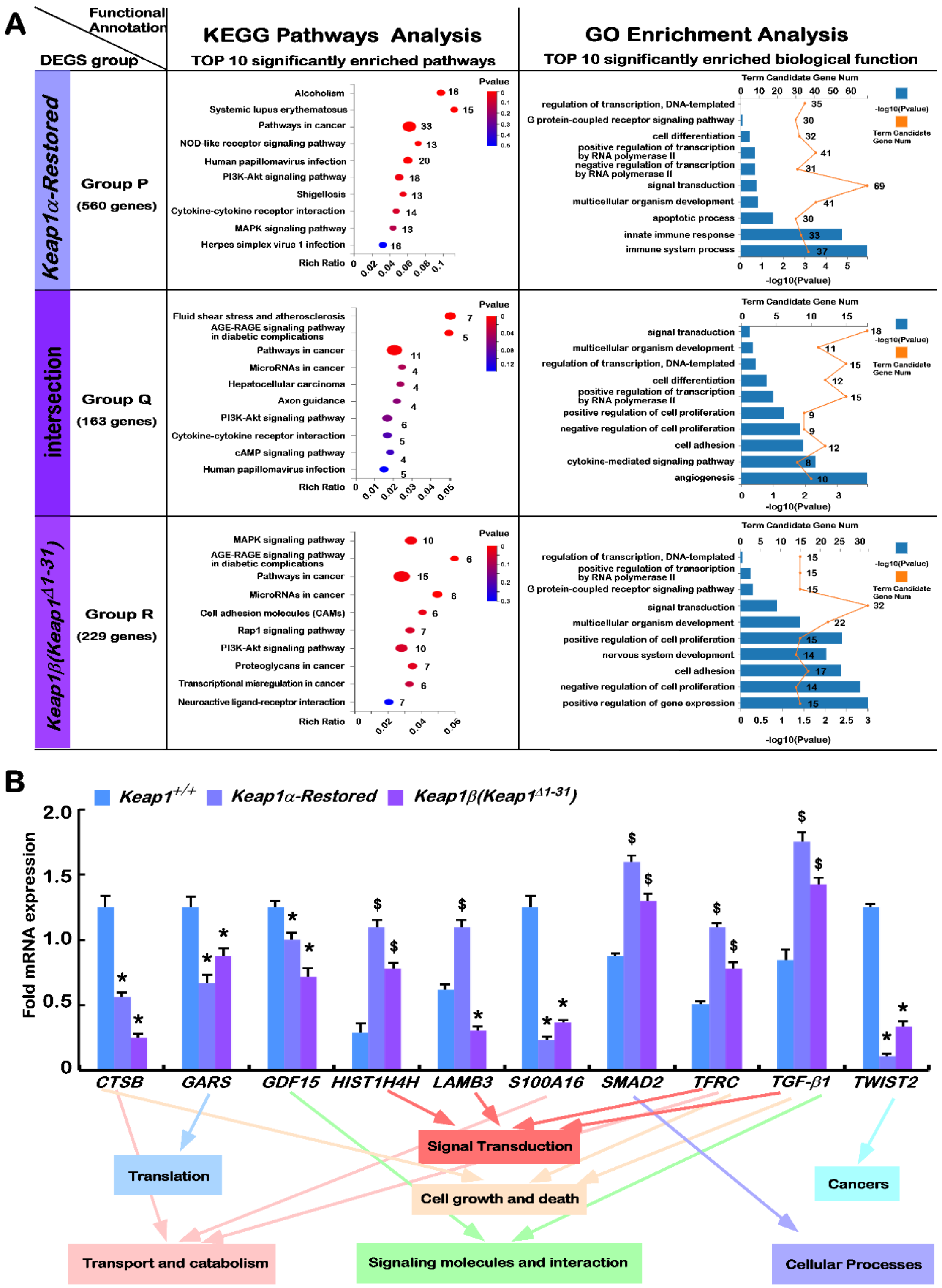


**Figure S8.** Functional annotation of specific or common DEGs in *Keap1α-Restored* and *Keap1β(Keap1$^{Δ1-31}$)* cells. **(A)** The top 10 of significant KEGG pathways terms and GO enriched by DEGs in Groups P, Q, and R were exhibited in scatterplots and histograms, respectively. Numbers indicate the number of genes. **(B)** Q-PCR was used to detect the mRNA expressions of *CTSB, GARS,GDF15,HIST1H4H,LAMB3, S100A16,SMAD2,TFRC, TGF-β1* and *TWIST2* in *Keap1$^{+/+}$*, *Keap1α-Restored* and *Keap1β(Keap1$^{Δ1-31}$)* cells. Putative functions of such target genes were mapped, as indicated by the histogram. The data are shown as mean± SEM(n=3X3, and "*" "$" indicates p≤0.01).

### Figure S9

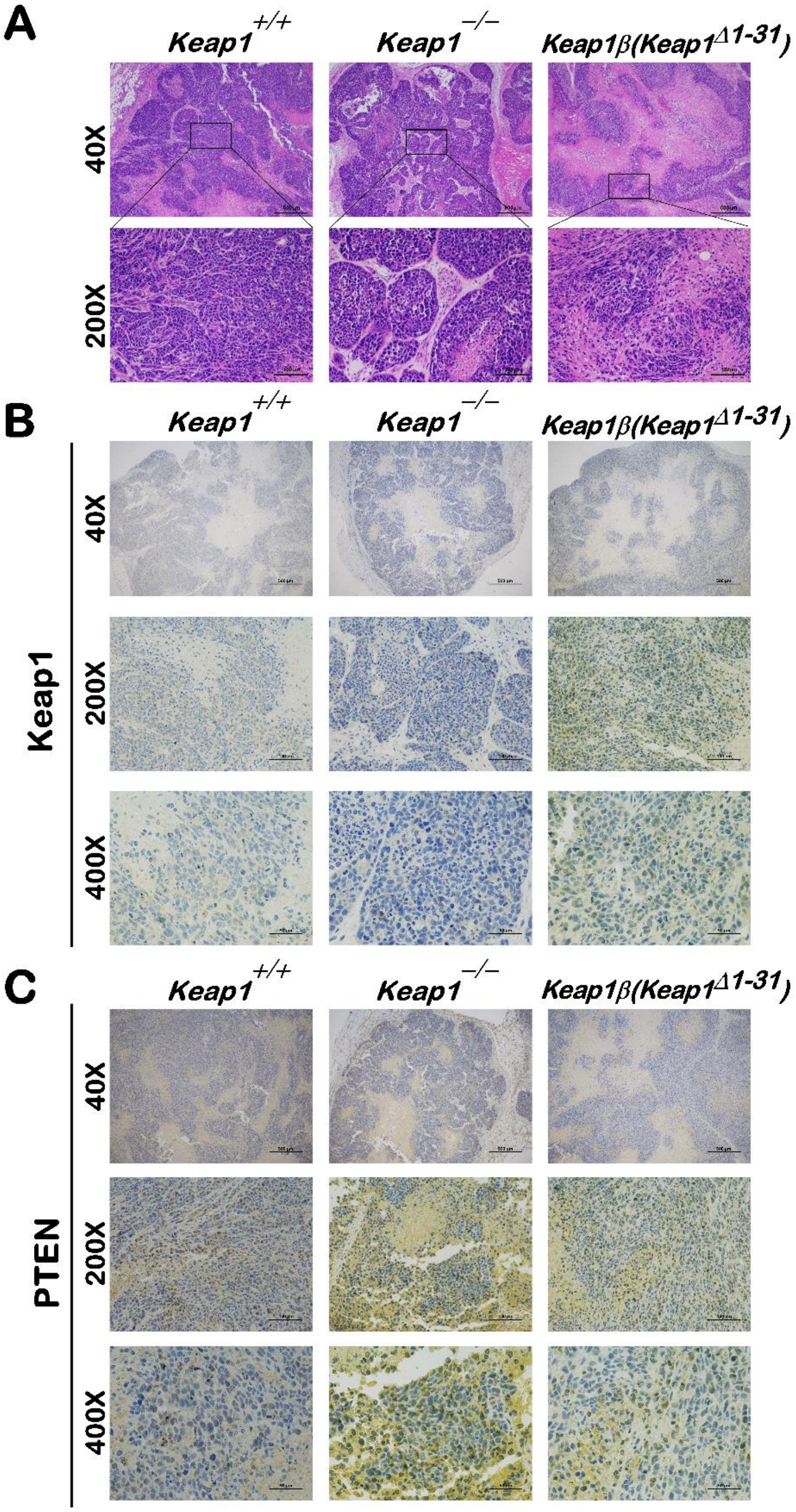


**Figure S9.** HE staining and immunohistochemistry of tumor tissue. Tumor from *Keap1*$^{+/+}$, *Keap1*$^{-/-}$ and *Keap1β(Keap1*$^{Δ1-31}$*)* cells grew under the skin of nude mice, and the tumors were removed to make tumor tissue slices. HE staining**(A)**, immunohistochemistry of Keap1**(B)** and PTEN**(C)** were performed, and photographs were taken. Pictures showed representative examples. H&E staining, Scale bars,100 μm; immunohistochemistry, Scale bars,500 μm,100μm and 50μm.

### Figure S10

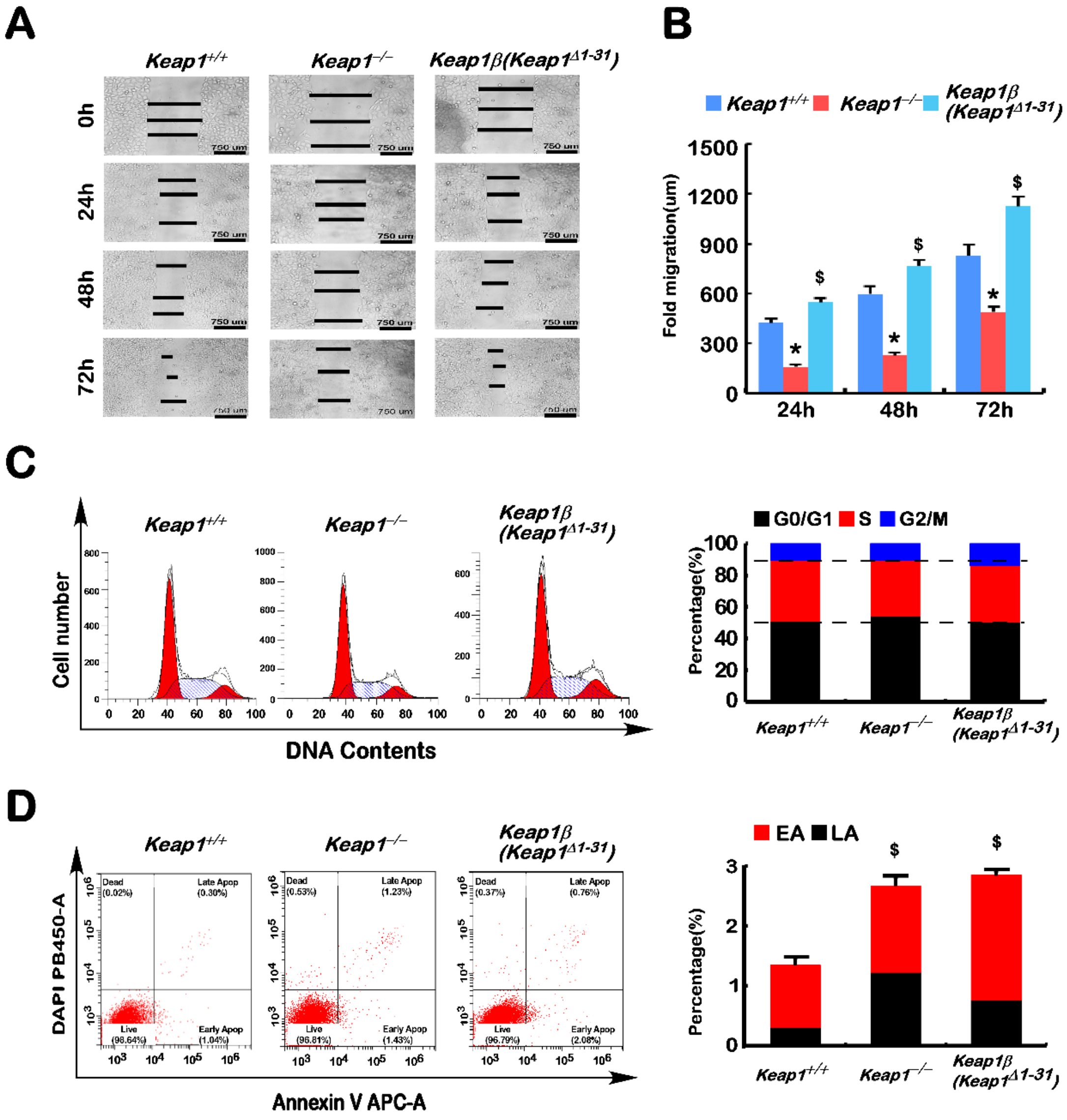


**Figure S10.** Analysis of wound healing, cell cycle and apoptosis. **(A)** Wound healing assay. *Keap1*$^{+/+}$, *Keap1*$^{-/-}$ and *Keap1β(Keap1*$^{Δ1-31}$*)* cells covered on the bottom of the small dish were scratched with the tip of 10 µL spear respectively, and the healing of each cell line was recorded at 0h, 24h, 48h and 72h by inverted microscope; Pictures showed representative examples. Scale bars,750 µm. **(B)** According to the scratch width in Figure A, the cell migration distance of each cell line was calculated at 24h, 48h and 72h, respectively. **(C)** Cell cycle detected by flow cytometry. Cell number distribution maps of *Keap1*$^{+/+}$, *Keap1*$^{-/-}$ and *Keap1β(Keap1*$^{Δ1-31}$*)* cell lines at each stage of the cell cycle were detected by flow cytometry. Figure C was used to calculate the percentage of *Keap1*$^{+/+}$, *Keap1*$^{-/-}$ and *Keap1β(Keap1*$^{Δ1-31}$*)* cells lines in G0/G1, S and G2/M, respectively. **(D)** Cell apoptosis was detected by flow cytometry. Cell apoptosis of *Keap1*$^{+/+}$, *Keap1*$^{-/-}$ and *Keap1β(Keap1*$^{Δ1-31}$*)* cell lines were detected by flow cytometry. The cells were divided into four zones: "Dead", "Late Apop", "Early Apop" and "Live". Figure D was used to calculate the percentage of apoptotic cells in *Keap1*$^{+/+}$, *Keap1*$^{-/-}$ and *Keap1β(Keap1*$^{Δ1-31}$*)* cells, with "LA" referring to Late Apop and "EA" to Early Apop. N≥3, and "*" "$" indicates p≤0.01.

### Figure S11

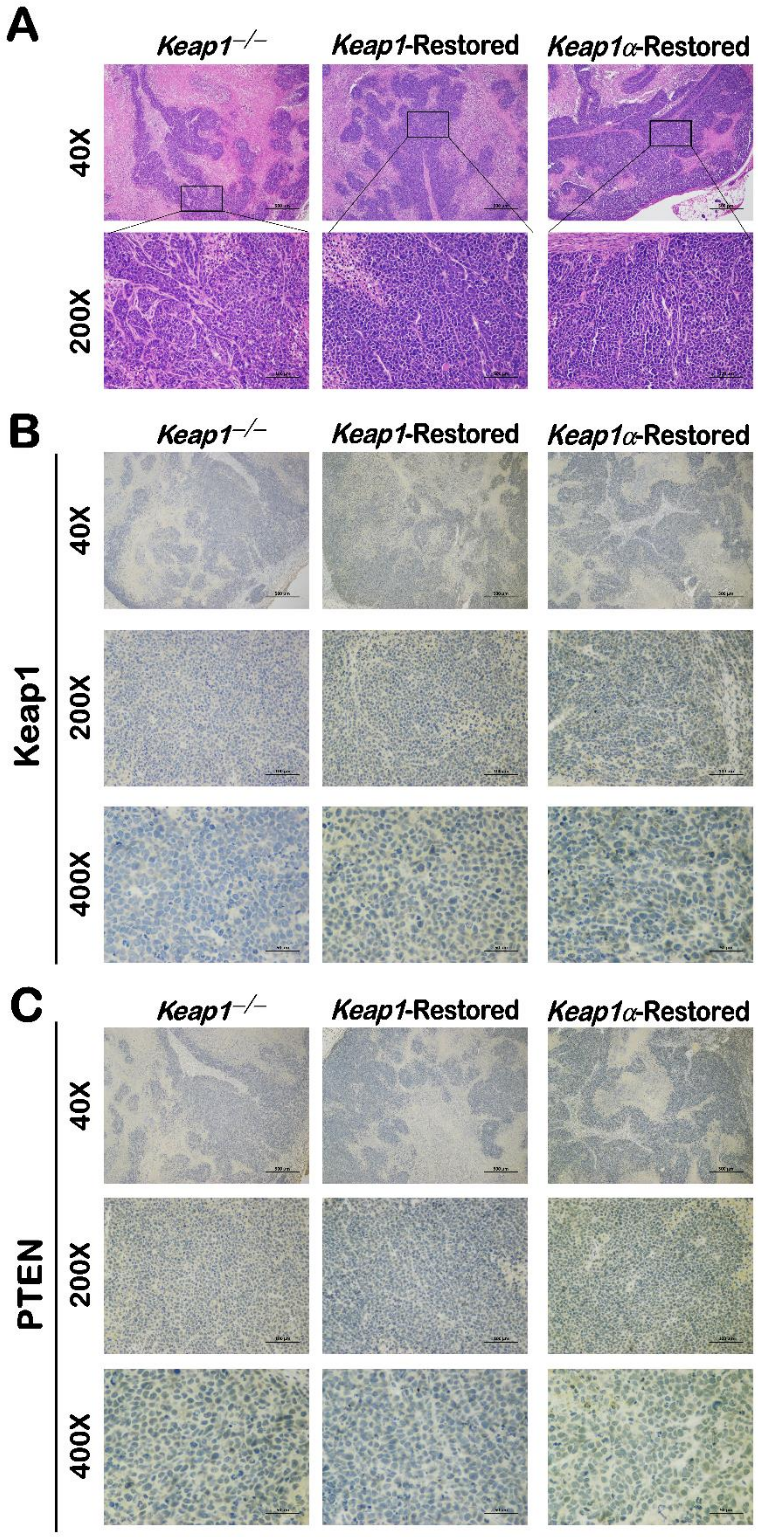


**Figure S11.** HE staining and immunohistochemistry of tumor tissue. Tumor from *Keap1*$^{-/-}$, *Keap1-Restored* and *Keap1α-Restored* cells grew under the skin of nude mice, and the tumors were removed to make tumor tissue slices. HE staining**(A)**, immunohistochemistry of Keap1**(B)** and PTEN**(C)** were performed, and photographs were taken. Pictures showed representative examples. H&E staining, Scale bars,100 μm; immunohistochemistry, Scale bars,500 μm,100μm and 50μm.

### Figure S12

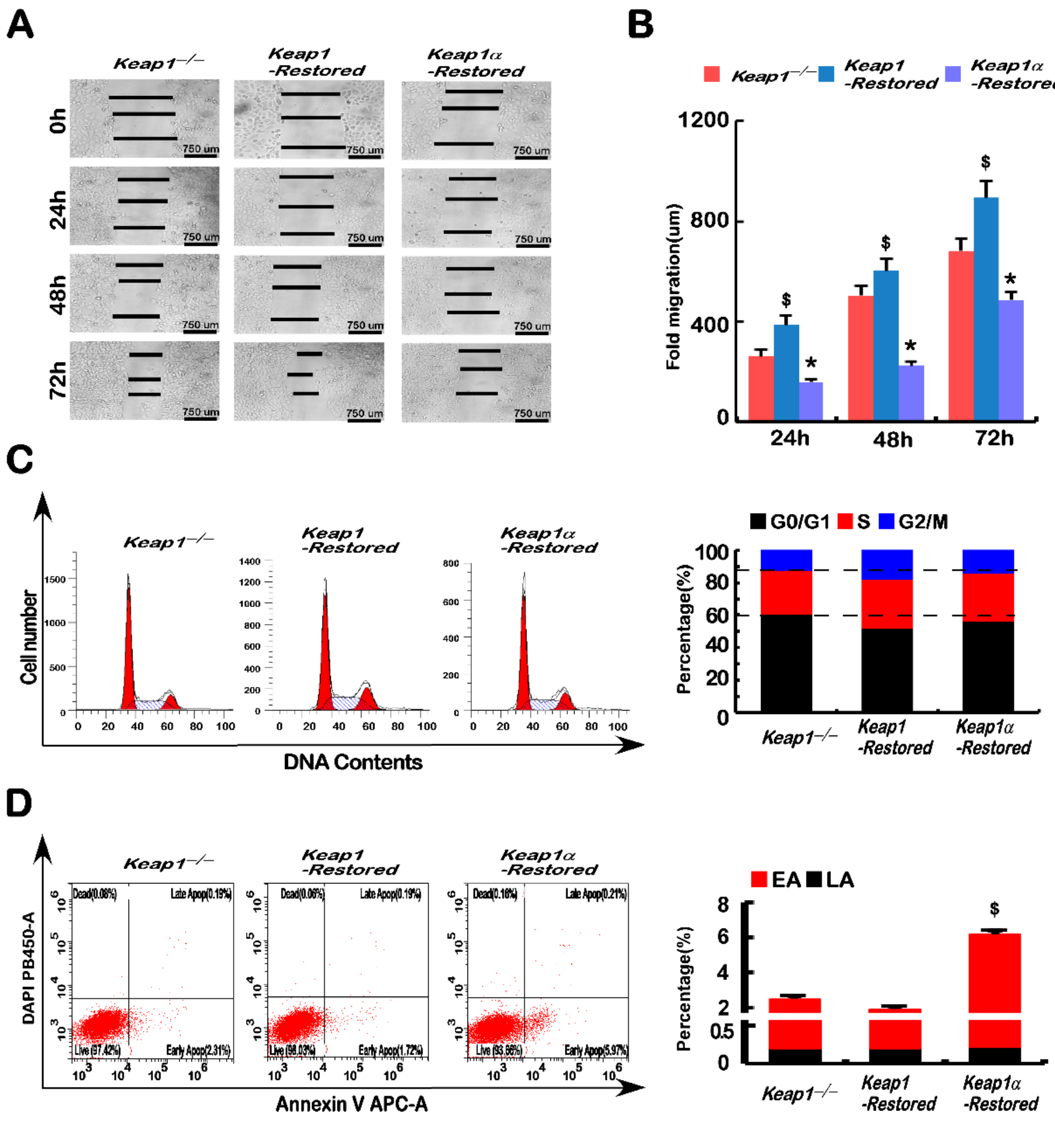


**Figure S12.** Analysis of wound healing, cell cycle and apoptosis. **(A)** Wound healing assay. *Keap1*$^{-/-}$, *Keap1-Restored* and *Keap1α-Restored* cells covered on the bottom of the small dish were scratched with the tip of 10 μL spear respectively, and the healing of each cell line was recorded at 0h, 24h, 48h and 72h by inverted microscope; Pictures showed representative examples. Scale bars,750μm. **(B)** According to the scratch width in figure A, the cell migration distance of each cell line was calculated at 24h, 48h and 72h, respectively. **(C)** Cell cycle detected by flow cytometry. Cell number distribution maps of *Keap1*$^{-/-}$, *Keap1-Restored* and *Keap1α-Restored* cell lines at each stage of the cell cycle were detected by flow cytometry. Figure C was used to calculate the percentage of *Keap1*$^{+/+}$, *Keap1*$^{-/-}$ and *Keap1β(Keap1*$^{Δ1\text{-}31}$*)* cells lines in G0/G1, S and G2/M, respectively. **(D)** Cell apoptosis was detected by flow cytometry. Cell apoptosis of *Keap1*$^{-/-}$, *Keap1-Restored* and *Keap1α-Restored* cell lines were detected by flow cytometry. The cells were divided into four zones: "Dead", "Late Apop", "Early Apop" and "Live". Figure D was used to calculate the percentage of apoptotic cells in *Keap1*$^{-/-}$, *Keap1-Restored* and *Keap1α-Restored* cells, with "LA" referring to Late Apop and "EA" to Early Apop. N≥3, and "*" "$" indicates p≤0.01.
